# Multimodal synaptomics reveals coordinated synaptic activity, protein synthesis and reorganization in NMDAR regulation

**DOI:** 10.1101/2024.10.28.620504

**Authors:** Reuven Falkovich, Sameer Aryal, Morgan Sheng, Mark Bathe

## Abstract

Complex neuronal circuit functions emerge from local, actively regulated synaptic protein levels that interplay with synaptic neurotransmission across heterogenous synapse populations. Understanding the mechanisms by which chemical and disease-associated genetic perturbations impact neuronal circuit functions requires simultaneous measurement of these factors with single-synapse resolution at population scale. Here, we combine in situ multimodal imaging of local mRNA translation, synaptic multiprotein composition, and synapse activity measured via calcium or glutamate fluxes, within the same spatially resolved synapses. We apply this approach of multimodal synapse profiling to study ketamine plasticity. Results map a causal network of NR2A-depletion-induced changes to synaptic scaffolding and receptor proteins, driven by synaptic activity and local mRNA translation, which translates to *Grin2a* models of schizophrenia *in vitro* and *in vivo*. Thus, multimodal synaptomics can reveal mechanistic neurobiology that underlies chemical and genetic perturbations within the context of scalable neuronal cultures, which can serve as models for human disease and therapeutic development.

## Introduction

Synapses shape neural circuits by integrating developmental, chemical, genetic, and activity cues and information into decisions of plasticity and metaplasticity. They achieve this via a complex interplay between dynamic levels of synaptic proteins that include receptor subunits^2^, scaffolding proteins^3,4^ and enzymes, mediated in part by local translation^5,6^ and signaling events such as phosphorylation^7,8^, calcium fluxes^9^, voltage changes, and neurotransmitter release. This complex synaptic protein-activity interaction system thereby plays a central role in the etiology and treatment of neurodevelopmental and psychiatric disorders^10–19^, as evidenced by the prevalence of synaptic proteins among disease-associated genes and psychiatric drug targets^18,19^.

Because the synaptic system is compartmentalized and locally regulated, numerous synapse-targeted methods have been developed to investigate their composition and disease-related changes, including synaptosome preparations^7,20–22^ or microdissection^1,22–24^. Combined with innovative labeling strategies, these approaches yield a wealth^22^ of detailed information on synaptic molecular composition. However, they lack the single-synapse-level resolution needed to resolve heterogeneity in both synapse composition and function, even amongst synapses of the same class^25^. Complemented by “synaptomic”^26–28^ approaches that map distributions of individual synapses in situ, using e.g. fluorescence or electron microscopy, additional valuable single-synapse ultrastructural and contextual information is provided, yet these are still typically limited either in their biochemical or activity information, or both. Thus, multimodal synaptomics that simultaneously accesses both molecular and activity information across individual synapses within intact neuronal systems could provide access to mechanistic, disease-relevant neural circuit functions that are otherwise impossible to measure. Such information could reveal how synaptic protein levels are either correlated or anti-correlated with synapse activities, for example, which is crucial to understanding the role of neuronal transmission in normal versus diseased states, and potentially also how to correct them using chemical compound screens. Yet, this information is currently impossible to access using separate protein labeling, neuronal activity, or bulk proteomic measurements, in vitro or in vivo.

Ideally, such a multimodal approach would incorporate information about synaptic mRNA and its translation events, layered on top of synaptic protein expression levels, and local synaptic activity. In particular, due to heterogenous subcellular localization in neurons, mRNA transcripts of many genes undergo a complex series of export, trafficking and retention^29–31^, protein binding^32–34^, modification^35,36^, translation^1,31^ and degradation that are locally regulated based on synaptic ion fluxes and protein activities^30^, which are affected via complex feedback mechanisms. Many synaptic proteins are synthesized on demand at the synapse where they are required^30,37,38^. This system plays a central role in normal as well as disease-associated or aberrant synaptic function^1,39^, for example in the case of fragile X syndrome^40,41^.

To realize *in situ*, simultaneous measurement of mRNA translation activity, multiprotein content, and calcium/glutamate spiking across millions of individual synapses, we combined multiplexed fluorescence imaging and Bayesian network analysis developed previously^42–44^ with *in situ*, single-synapse-resolution imaging-based sensors for calcium content^45^, glutamate release^46^, and gene-specific translation^47,48^ in neuronal culture. Previously, automated analysis yielded distributions of millions of synapses in high-dimensional protein space^42,44,49^, showing composition-defined subtypes of glutamatergic synapses, as well as complex synaptome-level phenotypes of activity perturbation^44^ or RNAi of genes associated with schizophrenia and autism spectrum disorder^49^. Bayesian network inference yielded network models of protein dependencies that predicted causal protein-protein effects inaccessible to conventional IF.

As a result, we observed function-composition dependencies that have previously only been inferred indirectly, such as between active versus silent synapses defined pre– and postsynaptically^50,51^. We demonstrated the value of this Multimodal-PRISM approach for in-depth exploration of concerted, interdependent subcellular mechanisms, focusing on plasticity induced by NMDAR antagonist psychedelics. We showed how this multimodal experiment can identify network mediators of treatment effects, yielding testable predictions of causal dependencies between plasticity in synaptic proteins, mRNA translation, and activity. Specifically, we identified processes of local activity-dependent and TrkB-involved synthesis and accumulation of synaptic proteins to maintain local NR2A levels in response to global perturbation. Finally, we illustrated how these observations might translate *in vivo*, identifying translation-dependent excitatory potentiation in an animal model of schizophrenia.

## Results

### CAT-PRISM allows mapping dependencies between synaptic proteins and calcium fluxes

To elucidate causal dependencies between synapse protein composition and synaptic activity, we sought to combine single-synapse activity trajectories with PRISM multiprotein imaging of the same synapses post-fixation. In our first reported method, termed Calcium Timelapse PRISM (“CAT-PRISM”, fig. 2a-d), we targeted calcium ion concentration dynamics, imaged using organic indicators. Since we anticipated slowly changing post-synaptic calcium levels to dominate the signal^52–56^, we imaged, in DIV 14 cultures, 30–60 minute-long timelapses at 3–6min per frame (Methods) to extract individual postsynaptic calcium traces over time. To enable assignment of synaptic calcium puncta to post-fixation synaptic PRISM signals, we used Hoechst staining present during both live and PRISM imaging, to align pre-to post-fixation images with a precision of ∼3–4 pixels, corresponding to ∼1 micron (Methods, fig. S1a).

In combined calcium-PRISM imaging at DIV 14 (Fig. 2a-d, Supp. Video 1), we identified that 35% of Synapsin1 puncta had corresponding calcium puncta from live-imaging. In order to visualize the distribution of calcium trace shapes, we performed dynamic time warping (DTW)^57^ (Methods) on mean-normalized traces, which groups traces of similar shapes closer together. The resulting distribution of calcium trace shapes was mostly one-dimensional (fig. S1b-c) with most traces monotonously increasing, decreasing or constant, and with additional differences corresponding to the abruptness of signal change and a small fraction of calcium-spiking puncta. Average synaptic calcium levels were consistently lower under APV NMDAR inhibition, whereas under TTX activity blockade they were initially higher than untreated synapses but subsequently lower over time (fig. 2b), matching the intuition that the indicator largely reports on activity-dependent postsynaptic calcium influx.

We combined DTW-analyzed calcium trace information with PRISM measurements of eight proteins— Synapsin1, Bassoon, VGLUT1, F-actin, CaMKIIα, GluR2, NR1, NR2B—in the same synapses (Methods) to yield a high-dimensional composition-activity distribution (fig. 2c) with clusters defined both by presence or absence of synaptic proteins and by calcium levels and trace shapes. To analyze this distribution in depth and identify direct protein correlates of synaptic activity, we extracted two main parameters describing calcium fluxes. The first was mean calcium level over time (“<Ca^2+^>”) and the second was the ratio of calcium level at the one-hour timepoint relative to the level at time zero (“ratio”). We then performed Bayesian Network analysis on the resulting 10-dimensional protein-activity distribution, as performed previously to summarize high-dimensional distributions as a graph of conditional probability relationships^49,58^, which presented testable predictions of causal dependencies between synaptic parameters^49^. The resulting Bayesian network (fig. 2d) identified NMDAR subunit levels (NR1 and NR2B) as directly correlated to average calcium levels, as anticipated. Calcium level ratio, on the other hand, was directly correlated to levels of CaMKIIα and negatively correlated with NR1 when controlling for all other proteins.

### GRL-PRISM reveals protein predictors of synaptic glutamate release behavior and correspondence between pre– and postsynaptically silent synapses

In our second reported method, termed Glutamate Release Loci PRISM (“GRL-PRISM”), we sought to incorporate a more direct indicator of synaptic activity by imaging single-synapse glutamate release events (“spikes”) using iGluSnFR3, a glutamate sensor with kinetics and signal-to-noise suitable for single synapse detection of minis and transmission events^46^. DIV 14 neurons transduced with iGluSnFR3 were imaged at 10– 20ms exposures, collecting 50–100 consecutive snapshots over 1–2s windows (fig. 2e, left. Supp. Video 2, Methods). Live images were again aligned to post-fixation PRISM images as described above. Spiking puncta were automatically identified using pixel-by-pixel spike detection on spatially smoothed images (fig. 2e, right, methods, fig. S1e) and assigned to post-fixation Synapsin1 puncta (fig. 2e, right) to enable combined analysis of ten protein measurements—the above set plus PSD95 and phospho-CaMKIIα—with spiking behavior of single synapses. Only a subset of Synapsin1 puncta are glutamatergic synapses, only a subset of these will have both active release machinery and iGluSnFR3, and only a subset of active synapses will be observed as firing during the short recording window employed. Thus, only ∼10% of Synapsin1 puncta had overlapping glutamate spiking puncta, although this number climbed to 20% when considering only synapses that contained all excitatory markers.

Mapping the distribution of synapses in 10-dimensional protein space (fig. 2f) revealed that synapses with at least one glutamate spike were drawn from a markedly different synaptic protein distribution than non-spiking ones. Specifically, they were enriched in regions 1 (all proteins present at high levels) and 2 (all proteins present at lower levels and GluR2 higher) at the expense of other regions, which lacked GluR2 and other protein combinations (fig. 2f, right). Chemically-induced LTP (20uM Bicuculline + 200uM glycine)^59^ increased both the frequency and intensity of Synapsin1-associated optical spikes over time over control (BC alone) cultures (fig. 2g).

We then sought to perform a more systematic analysis of the protein correlates of glutamate spiking, and specifically of three aspects of activity: (i) existence of a spike, (ii) overall intensity of spikes, and (iii) increase in spike intensity, which indicates synaptic potentiation. We performed a logistic regression for measure (i) and linear regressions for measures (ii) and (iii) against the ten measured proteins (fig. 2h). We observed that Synapsin1 and VGLUT1 were strong predictors of spiking probability and intensity. The coefficients of Synapsin1 may be partially artifactual, however, because Synapsin1 puncta were used to define synaptic regions in images with protein overlap used to assign spiking analysis puncta to synapses, which may be more likely with larger puncta.

Phosphorylation of CaMKIIα was also predictive of spiking probability and intensity, consistent with its dependence on calcium influx from synapse activity and its contribution to LTP^8,60,61^. We were surprised to observe strong negative coefficients of NR1 in spiking probability and intensity. When simultaneously measuring NR1 (which accounts for both NR2A and NR2B-containing NMDARs) and NR2B, the remaining variance in NR1 was at least in part due to NR2A levels. This indicates a negative dependence of synapse glutamate activity on NR2A with constant NR2B and GluR2, which we explored further in figures 4-6. Regression coefficients of spiking potentiation indicated negative and positive coefficients for F-actin and PSD95, respectively, as well as a negative coefficient for NR2B (fig. 2h). A hypothetical interpretation of this observation is that larger dendritic spines with smaller amounts of PSD95 and NR2B-containing NMDA receptors indicates a synapse in the process of being reduced or eliminated prior to the spine shrinking as well, whereas larger amounts of PSD95 with smaller dendritic spines may indicate a synapse in the process of potentiating^62^.

Finally, GluR2 had the second-highest coefficient for spiking probability but no contribution to spike intensity, matching our observation of spiking synapses originating almost exclusively from GluR2-containing clusters. This is consistent with a model of binary categories of silent and active synapses and presents a direct observation that pre-synaptic classification of these categories (defined by vesicle release)^50^ matches the post-synaptic classification (defined by AMPAR presence)^51^, thereby reinforcing the importance of combining activity and protein measurements within the same synapses.

### Integration of *in situ* mRNA and translation labeling into Multimodal-PRISM

In order to incorporate *in situ* measurement of gene-specific local synaptic translation into PRISM, we developed RIBOmap-PRISM (Methods), an adaptation of ribosome-bound mRNA mapping (RIBOmap)^47^ to PRISM probe-exchange multiplexing. RIBOmap identifies putative mRNA translation events *in situ* using coincidence detection between mRNA transcript-specific primer and padlock probe pairs and a probe against the 18S ribosomal RNA. The combination of all three generates a single-stranded RCA amplicon when spatially co-localized. In RIBOmap-PRISM, the resulting amplicon presents a docking strand, which can then be imaged in a standard PRISM protocol alongside PRISM barcoded antibodies (fig. 1b, 3a). We also adapted spatially-resolved transcript amplicon readout mapping (STARmap)^48^, to identify mRNA transcripts regardless of ribosome association, into STARmap-PRISM in a similar fashion (fig. S2a).

**Figure 1:**
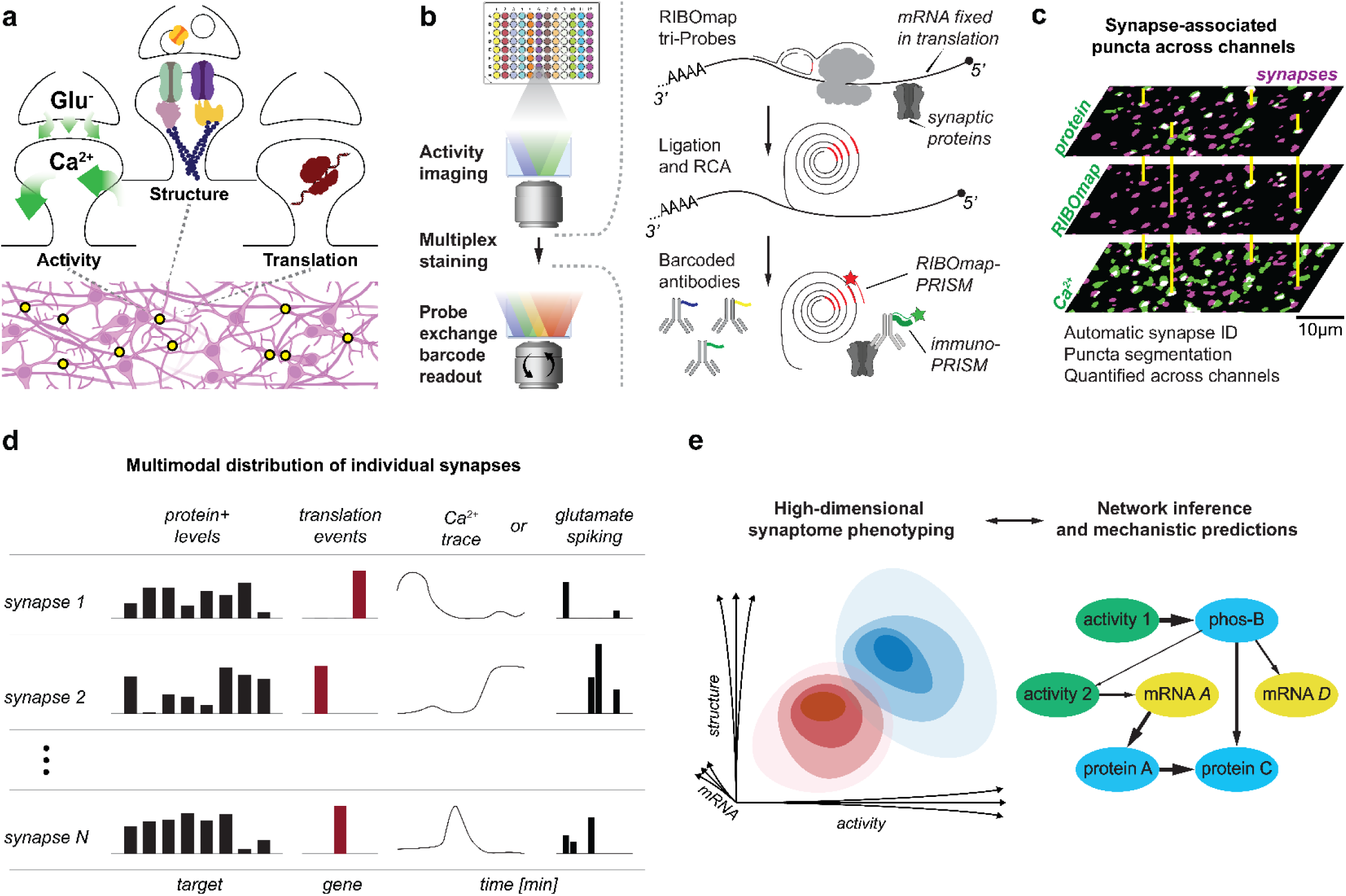
Summary of the Multimodal PRISM approach. A) Each synapse from a large heterogeneous population of a neuronal system is a compartmentalized interacting system between activity represented by ion fluxes, protein composition and protein synthesis. B) Tandem live imaging followed by a combination of in-situ methods that associate DNA barcodes with proteins and molecular events such as mRNA translation. This is followed by multiplexed PRISM barcode readout using fluorescent LNA probe exchange. C) The same field of view imaged across many rounds exhibits puncta associated with different synaptic parameters, which are automatically identified, associated with synapses, and quantified. D) The resulting data is a high-dimensional distribution of individual synapses with information about multiple proteins and phosphoproteins, mRNA translation, and synaptic activity, for each. E) This in turn enables identification of subtle gene-or drug-induced phenotypes in the synapse distribution not visible to bulk methods (left), as well as generating models of causal dependencies between synaptic parameters and changes to them (right).

This approach generated discrete puncta corresponding to translation events of *Actb* (β-actin), *Dlg4* (PSD95), *Camk2a* (CaMKIIα) and *Ntrk2* (TrkB) (fig. 3a). Identifying synapses, dendrites, and soma using Synapsin1 and MAP2 staining in the same sample (Methods) allowed us to assign RIBOmap-PRISM puncta to these neuronal regions (fig. 3b) and compare them with other measurements of subcellular transcript localization and translation, specifically somatic versus neuropil ribosomal profiling (Ribo-Seq) from microdissected tissue^1^. Counting puncta enrichment in dendrites versus soma, we observed that these four genes followed similar patterns as neuropil/soma enrichment levels measured by Ribo-Seq (fig. 3c), despite differences in model systems and measurement approaches. Estimates (Methods) of the number of identified translation events per cell in RIBOmap-PRISM matched those from Ribo-Seq (fig. S2b). Both synaptic and overall RIBOmap-PRISM puncta counts of *Dlg4* and *Grin2a* were reduced with respective siRNA-mediated knockdown (fig. 3d and S2c). Dendrite– and synapse-localized puncta of *Dlg4* were more affected by its siRNA (∼47% reduction) than somatic puncta (∼19% reduction) with a corresponding change in neurite enrichment (log-fold of 1.02±0.24 in NonT-treated cultures vs 0.52±0.17 in *Dlg4*-treated cultures), which agrees with a steady state in which transcripts are constantly produced but a fraction is degraded via RNAi before they can reach distal regions. Finally, we tested a gene (*cox1*) that is known to be translated inside mitochondria^63^ and observed that RIBOmap-PRISM puncta of this transcript colocalized with mitochondria identified using CoxIV staining (fig. S2d-f).

**Figure 2:**
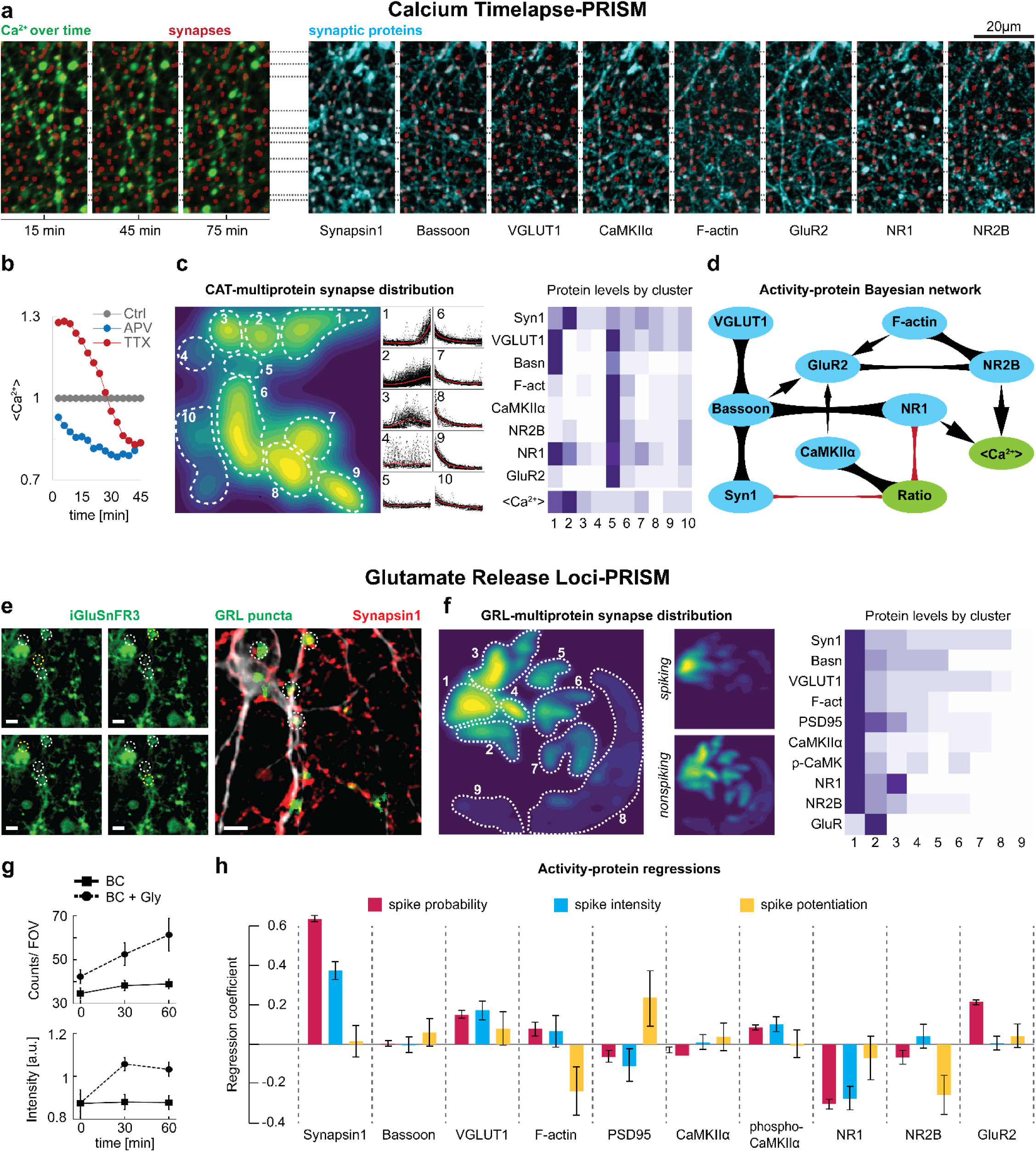
Live-PRISM. A-D: CAT-PRISM. A) Aligned images of the same field of view as in fig. 1c during live imaging of a calcium indicator before fixation (green), and over sequential imaging rounds of different synaptic proteins (cyan), overlaid with synaptic puncta (red) defined by Synapsin1. B) Average calcium signal in synapse-associated puncta in cultures treated with TTX or APV, relative to untreated. C) Left: Density contour plot of a UMAP projection of synapse distributions in protein levels, average calcium levels, and calcium trace shapes, with manually identified clusters. Middle: All (gray) and average (red) calcium traces in each of the clusters 1-10 of (C). Right: Average protein and calcium levels in each of the clusters. Columns are clusters 1-10 in order. D) Bayesian network connecting eight protein levels and two activity measures. Arrow thickness denotes edge strengths, defined as parent-controlled correlations (see Methods). Black arrows indicate positive correlations, red indicate negative. E-H: GRL-PRISM. E) Left: Four frames of the same FOV, each with a different region spiking relative to the other three regions (circled). Right: The same FOV, overlaying spike detection output (green) with post-fixation Synapsin1 staining (red) and MAP2 (white). Scale bar is 10um. F) Left: Distribution of synapses in protein space (density contour plot of UMAP projection). Small: sub-distributions of synapses with and without a detected spike. Right: Relative average protein levels of regions delineated in (F). G) Spike counts and average intensity at different times during treatment with chemical LTP (Bicuculline + glycine, compared to Bicuculline baseline). Error bars are mean ± s.e.m across wells. H) Regression coefficients of spiking paramaters against the nine measured proteins. Red: logistic regression for spiking binary outcome. Blue: Linear regression for spike intensity among spiking synapses. Yellow: Difference between spike intensity at 1 hour and at start. Error bars are S.E. of the regression.

**Figure 3:**
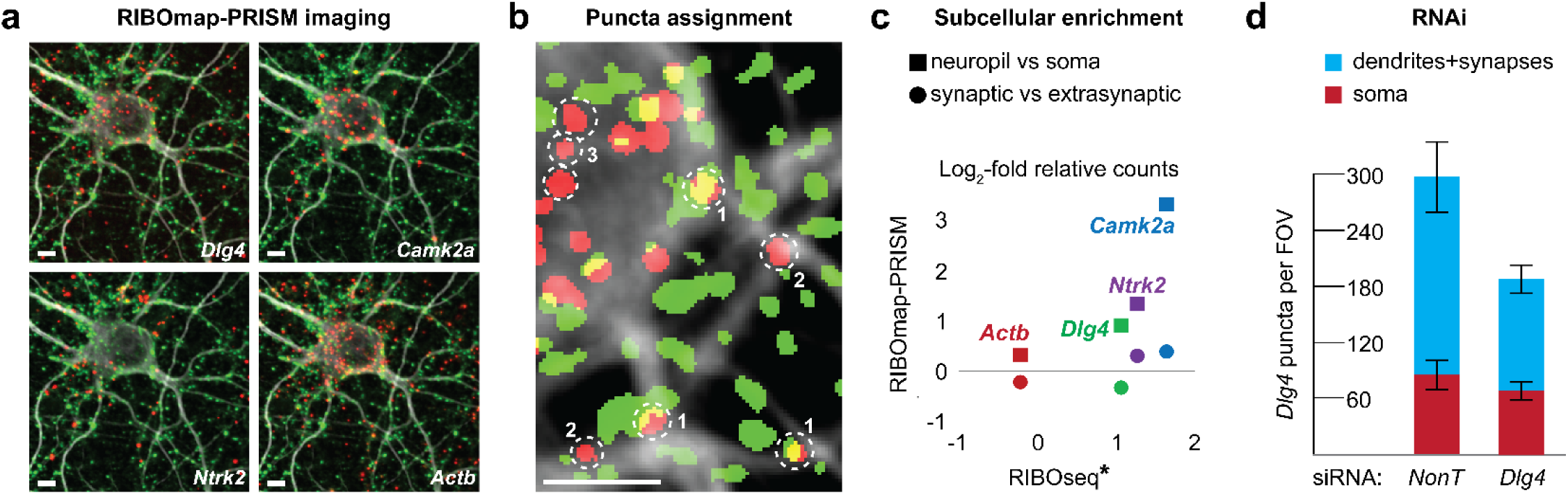
RIBOmap-PRISM. A) Sequential images of RIBOmap-PRISM (red) of four different genes, co-stained with MAP2 (white) and Synapsin1 (green) captured over four sequential PRISM probe exchange rounds. B) Automatically segmented puncta of *Dlg4* RIBOmap-PRISM (red) and synapses (green) overlaid over MAP2 staining (white), showing translation events that are synaptic (1), neuropil-localized extrasynaptic (2) and soma-localized (3). Scale bar in A and B is 5µm. C) Ratio of RIBOmap-PRISM puncta associated with different subcellular structures (soma/neuropils, and synaptic vs neuropil extrasynaptic), presented in log2 scale. Y-axis: Ratios obtained from RIBOmap-PRISM, X-axis: ratios obtained by Glock et al.^1^ using microdissection and RIBOseq, which only provides information on neuropil vs soma ratios. D) Count of somatic (red) and dendritic+synaptic (blue) puncta of *Dlg4* per field of view after 8 days of treatment with nontargeting or *Dlg4* siRNA. Error bars are mean ± s.e.m. across wells.

To test whether amplicons generated by RIBOmap-PRISM crowded one another or affected accessibility of protein antigens in tight synapses, we examined *Dlg4* RIBOmap-PRISM puncta intensity, shape, Synapsin1 signal and probability of *Ntrk2* puncta colocalization in synapses containing *Dlg4* RIBOmap-PRISM puncta that underwent different durations of amplification. Neither of the latter were affected despite a ∼7-fold increase in RCA amplicon size, as indicated by *Dlg4* RIBOmap-PRISM integrated fluorescence signal (fig. S2g). We also observed (fig. S2h-i) that even following low amplification durations the distribution of RIBOmap-PRISM intensities in synapses was approximately exponential with a single mode, consistent with the expected distribution of single amplicon sizes^64^, which indicated that very few synapses, if any, contained more than one puncta for this gene. This is consistent with observations of few polysomes and ∼2–3 translation events (of any transcript) per synapse^37^ at the moment of fixation. Thus, for all subsequent analyses, we treated the RIBOmap-PRISM signal as a binary variable indicating the presence or absence of a single ribosome-associated mRNA for a specific gene.

### Measuring concerted synaptic activity-translation-multiprotein changes under NMDAR antagonism

Ketamine is an NMDAR antagonist which can act as both a potent dissociative psychedelic and an antidepressant with both rapid onset and sustained action, with a wide range of proposed mechanisms^65,66^ to connect these cognitive effects to the molecular changes it elicits. We sought to apply Multimodal PRISM to explore concerted and interdependent changes to synaptic activity and structure induced by ketamine’s NMDAR antagonist action at the single-synapse and population level, building on prior observations that chronic Grin2a knockdown and APV NMDAR blockade elicit a potentiation in excitatory synapse density and protein levels^49^. We performed live glutamate imaging on DIV 14 neuronal cultures treated with Ketamine or PCP in either an acute (immediately before live imaging) or chronic manner (DIV 6-14, removed before live imaging) (fig. 4a-b). These cultures were then fixed and PRISM-stained for a set of proteins and phosphoproteins as well as RIBOmap-PRISM of *Camk2a* (fig. S3a). These included Synapsin1, presynaptic proteins VGLUT1 and Bassoon, AMPAR and NMDAR subunits GluR2, NR2A and NR2B, NDUFA9 to mark synapse-associated mitochondria, and phosphorylation levels of ERK and S6RP, marking activation of the TrkB signaling pathway to local synaptic translation, thought to be implicated in the long-term nature of its antidepressant effects^67,68^.

Cultures chronically treated with ketamine showed increases in the number of glutamate release spikes, both immediately and 1.5 hours after drug removal (fig. 4c and S3b), alongside a decrease in average spike intensity but stronger intensity *potentiation* among synapses that spiked at both timepoints. They also showed increases in the synaptic levels of all measured proteins including phospho-ERK and phospho-S6RP (fig. 4d), a fourfold increase in the fraction of synapses translating *Camk2a* (fig. 4e), and an increase in the fraction of VGLUT1-positive synapses (fig. 4f). Interestingly, acutely treated cultures exhibited opposite or no effects, except for similar effects on spike intensities. These may indicate that the drug simultaneously weakens synapse strength in culture and primes them for stronger potentiation once the drug is removed, and that the chronic treatment system may act as a more representative in vitro model, despite the discrepancy in timing from in vivo observations^68^, which we discuss in the next section.

PCP exhibited directionally similar effects (fig. S3b-c), with a prominent difference of a three-fold reduction in synaptic NR2A under acute treatment. These observations were congruent with our previous report^49^ that short-term treatment with NMDAR antagonist APV elicited a decrease in synaptic NMDA receptors, while under long chronic treatment this effect disappeared, and cultures exhibited an increase in excitatory synapse density and levels of excitatory postsynaptic proteins. Here, chronic treatment with APV similarly elicited a threefold stronger increase in glutamate spiking events over time after drug removal (fig. S3b, right).

We thus decided to use APV as a model for an in-depth exploration and mechanistic analysis of synaptic activity, translation and structural changes associated with chronic NMDAR inhibition. Since effects of activity on postsynaptic protein rearrangement are largely mediated by calcium flux, we performed CAT-PRISM to measure calcium changes, translation of three genes, as well as puromycin incorporation to ^69,70^ report on overall translation (fig. S4a), and eight proteins in the same synapses (fig 5a-b). We observed an increase in presynaptic (Bassoon) as well as postsynaptic scaffolding (CaMKIIα, PSD95, F-actin) and glutamate receptor (GluR2, NR2B) proteins. We also observed a decrease in starting calcium levels, an increase in calcium end-to-start ratios, and an increase in excitatory synapse density, as expected from the changes in spiking intensity. These changes accumulated over treatment time (fig. S4b-f), with synaptic NR2A levels lowered at short treatments before returning to baseline, as we previously observed.

**Figure 4:**
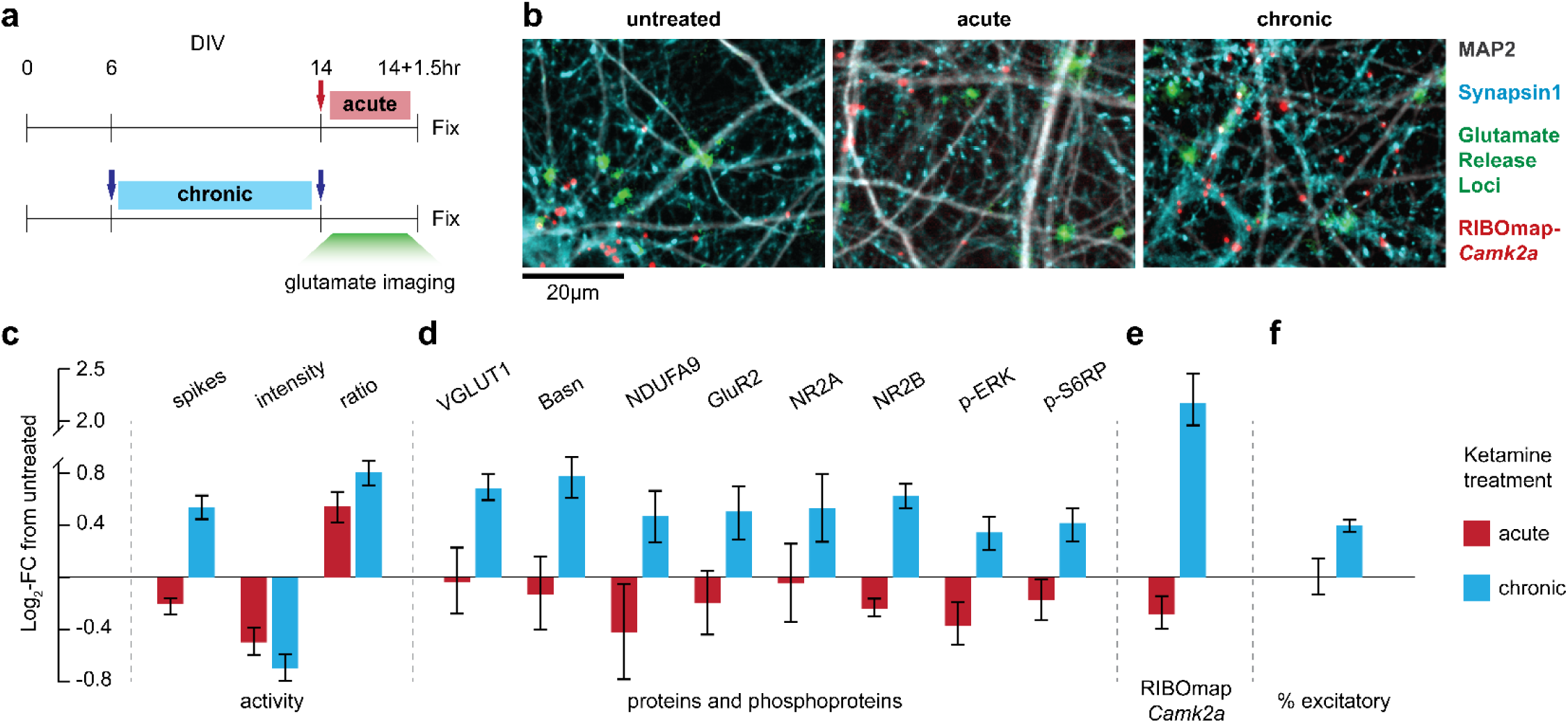
Ketamine-induced plasticity. A) Treatment schematic. Arrows indicate when ketamine or PCP were added and/or removed in each regime. B) Aligned images of glutamate spike detection (Methods) from imaging pre-fixation (green) with MAP2 (white) and Synapsin1 (red) staining post-fixation of the same FOV. C-F) Changes in synaptic parameters in acute– and chronic-treated cultures relative to untreated. C) Overall number of glutamate spikes (combined early and late imaging), average spike intensity of synapses that spiked in early imaging, and ratio of late to early intensity of synapses that spiked in both time points. D) Average synaptic levels of 6 proteins and two phospho-proteins, imaged with Immuno-PRISM. E) Fraction of synapses containing *Camk2a* RIBOmap puncta. F) Fraction of synapses positive for VGLUT1. C-F error bars are means ± s.e.m. across wells.

**Figure 5:**
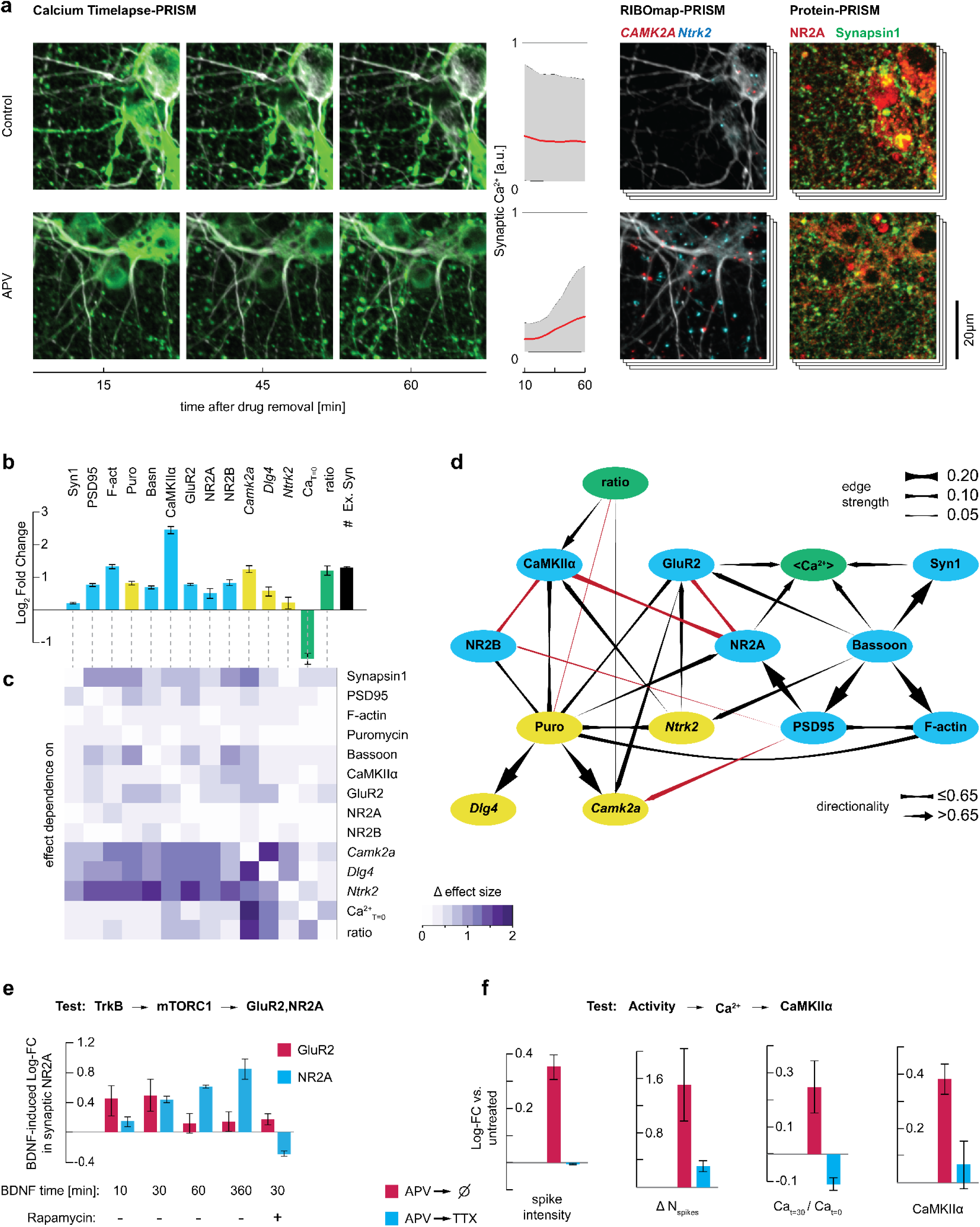
In-depth investigation of causal dependencies in the multimodal synaptic response. A) Left: Sequential images of the same fields of view with three timepoints of Fluo-8FF. Middle: Synaptic calcium traces from the Fluo-8FF imaging. Average (red) and 5-95^th^ percentile (gray). Top and bottom graphs on the same scale. Right: one of the RIBOmap-PRISM rounds, and one of the Protein-PRISM rounds, of the same field of view imaged left. Calcium and RIBOmap-PRISM images overlaid with post-fixation MAP2 staining (white). Top row from untreated culture, bottom row for culture after chronic NMDAR blockade with APV. B) Changes to synaptic proteins (blue), translation indicators (red), and activity parameters (yellow) as well as excitatory synapse density (black) in chronically APV-treated cultures, relative to untreated. C) Dependency of the effects of APV treatment on each synaptic parameters (columns) when controlling for each of the other parameters (rows). Heatmap generates the difference between the effect size in synapses at the top quartile of the controlling parameter, versus synapses at the bottom quartile, and this is normalized to overall effect size (see Methods). D) Inferred combined Bayesian network of eight protein, four translation and two activity variables, in untreated cultures. Black arrows and lines denote positive edges, red negative. This network is partial contains only some of the edges, for a full network shown as a heatmap see figure S5b. E) One prediction from C and D, testing by measuring synaptic NR2A after treatment with BDNF or BDNF+rapamycin. F) A second prediction from C and D, tested by measuring synapses treated with TTX after removal of APV, which stops the increased activity of APV-potentiated cultures. Error bars are mean ± s.e.m. across wells.

These accompanied an increase in local protein synthesis, both overall (by puromycin) and specifically in fractions of synapses containing *Camk2a*, *Dlg4* (PSD95), and *Ntrk2* (TrkB) RIBOmap-PRISM puncta. The latter possibly being replacement of internalized TrkB units after activation^71–73^. When counting overall RIBOmap– and STARmap-PRISM puncta, we observed strong increases in neurite-localized RIBOmap-PRISM, with considerably smaller increases and even decreases in soma-localized RIBOmap-PRISM and overall STARmap-PRISM, supporting our interpretation of a subcellularly targeted translation-dependent response. Unlike the three genes above, transcription and translation of non-neuropil-targeted Actb were unaffected (fig. S4h).

Interestingly, chronic treatment with NR2A-specific antagonist MPX-004 yielded no or opposite effects across most proteins, RIBOmap and activity measures, and a transient increase in NR2A (fig. S4c, S4j and S5a), while the NR2B-specific antagonist Traxoprodil-CP101601 yielded responses similar to APV but weaker (fig. S4j). Furthermore, we observed that APV treatment, but not MPX, reduced overall RIBOmap-PRISM signal of Grin2a (fig. S4g). These observations, combined with our previous observations of similar excitatory synaptic protein potentiation in siRNA-mediated Grin2a knockdown^49^, indicated that the observed synaptic changes were likely downstream of global reduction in NR2A expression caused by pan-NMDAR or NR2B antagonism, rather than a response to perturbed local activity of NR2A. Moreover, given the transient change in synaptic NR2A, the opposite effects in synaptic NR2A and other proteins under MPX-004, and our observations that Grin2a RNAi knockdown reduced RIBOmap signal but not synaptic Grin2a levels (fig. S2f and S4k), these were possibly part of a compensatory process that maintains steady local synaptic NR2A levels in the face of changes to global supply.

### Multimodal Bayesian network and conditional effects analysis generates testable predictions of causal dependencies between synaptic processes

To investigate causal dependencies between changes to synaptic proteins, translation, and activity, we constructed a map of the effect of each synaptic measure on the APV-elicited phenotype in each other measure (fig. 5b-c, Methods). We also constructed a Bayesian network connecting eight protein measures, four local translation measures, and two activity measures (fig. 5d). Together, the map of conditional effects and Bayesian network yielded several observations that translate to testable predictions of dependencies between synaptic changes.

First, we hypothesized that, if the potentiation in excitatory synaptic proteins involves TrkB-driven local synaptic translation (as evidenced by the edges connecting puromycin and RM-*Ntrk2* to other synaptic proteins), and if that potentiation is tied to maintaining synaptic NR2A levels, then BDNF stimulation should increase synaptic NR2A in an mTORC1-dependent manner. We thus tested synaptic GluR2 and NR2A levels at different time points after adding BDNF, with and without rapamycin (fig. 5e). We observed that NR2A levels accumulated with TrkB activation, and that this was abolished by mTORC1 inhibition.

Second, we observed that the increase in local translation of *Camk2a* depended on changes to synaptic activity (fig. 5c), consistent with the Bayesian network edge from Ca2+ ratio to CaMKIIα. We thus hypothesized that the local translation-driven increase in CaMKIIα depends on changes to synaptic activity induced by chronic APV treatment. To test this, we tested APV-induced changes when culture activity was blocked by TTX during the recovery period (during live imaging) (fig. 5f). Indeed, blocking activity abolished the APV-induced potentiation in spike intensity and number, calcium accumulation, and CaMKIIα, but did not affect the changes to excitatory synapse density or starting calcium (fig. S5d).

Further mechanistic predictions can be made with additional observations in the Bayesian network and its changes; for example, the most strongly changed network edges were those surrounding CaMKIIα (fig. S5b-c), indicating that some unmeasured proteins or processes were involved in its activity-dependent translation and accumulation at the synapse. Finally, the negative edges observed between NR2A and GluR2 or CaMKIIα in the Bayesian network (fig. 5d) were consistent with the negative correlation of NR1 to Ca increase (fig. 2f) and contribution to synaptic spiking and intensity (fig. 2k) when controlling for NR2B, consistent with a scenario in which NR2A depletion triggered increases in these proteins, and may indicate that these proteins’ increase is incidental rather than necessary for NR2A maintenance.

### NR2A depletion phenotype extends to Grin2a KO models of schizophrenia in vitro and in vivo

The causal inferences above were made for synapses in an in vitro neuronal culture. However, they describe fundamental aspects of glutamatergic synapse organization that may be speculated to operate across organisms, as well as in vivo. In addition, these observations indicate that the translation-dependent potentiation processes are downstream of global NR2A depletion. Thus, we predicted that we might observe corresponding processes in *Grin2a* knockout mice, a genetic model for schizophrenia with a high penetrance gene and considerable face validity^74^.

Towards this end, we observed increases in excitatory synaptic proteins, puromycin incorporation, and synapse density in cultures from *Grin2a* +/-pups compared to litter-matched *Grin2a* +/+ controls (fig. 6a-b). The multiprotein Bayesian network again showed a strong direct relationship between NR2A and local synaptic translation indicated by puromycin (fig. 6b). Finally, measuring synaptic proteins in postsynaptic density (PSD) fractions in these mice in vivo, as well as their corresponding mRNA in bulk RNA-seq from cortical regions (fig. 6c), we observed increases in excitatory synaptic proteins without corresponding increases in transcript levels, consistent with translation-dependent potentiation.

**Figure 5:**
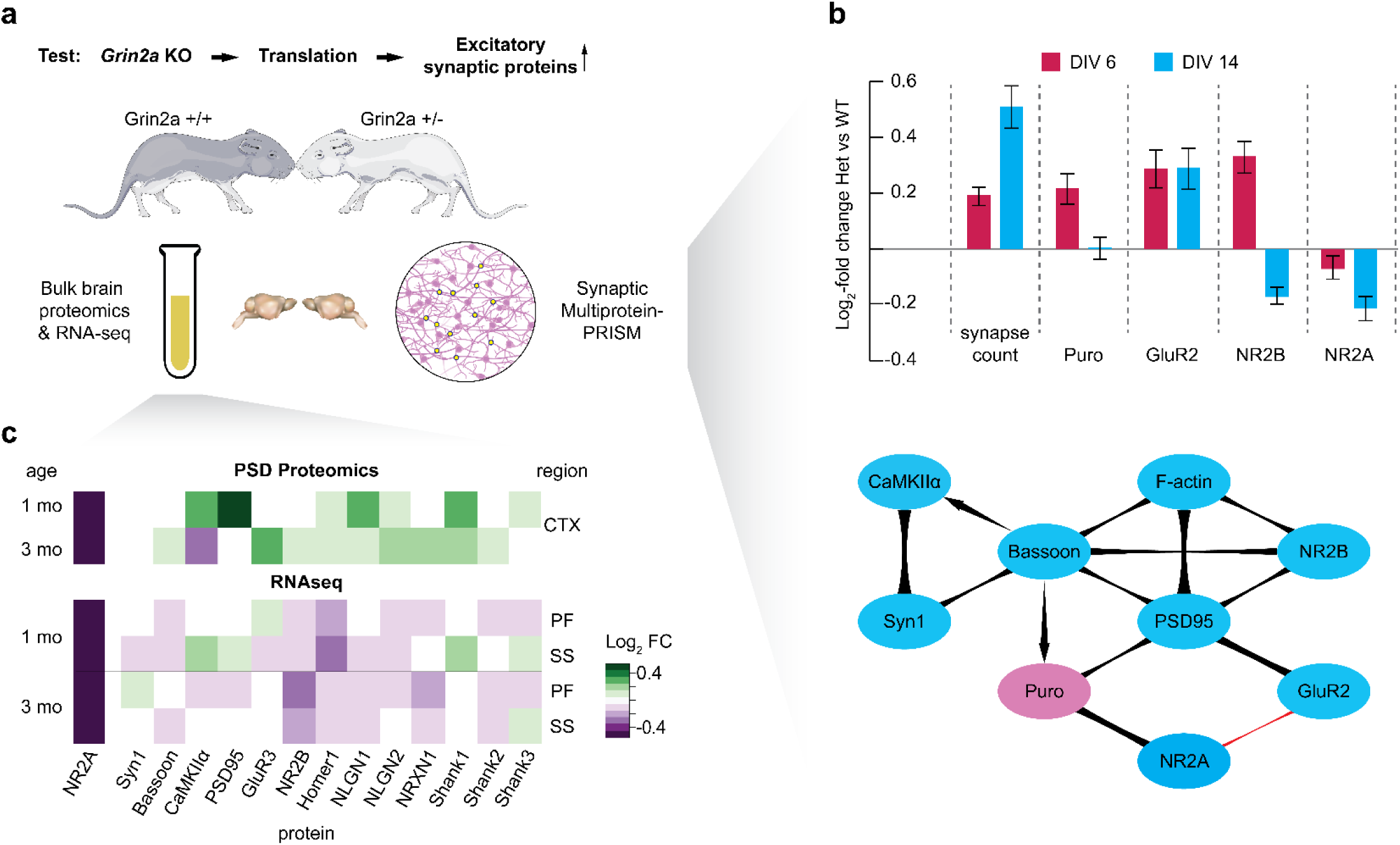
Translation of NMDAR antagonism observations to *Grin2a* KO mice in vitro and in vivo. A) Prediction and schematic. B) Single-synapse, multiprotein measurements of mouse *Grin2a* +/-neurons in vitro. Top: Changes to average excitatory synapse density, puromycin incorporation, and receptor levels, relative to neurons from litter-matched *Grin2a* +/+ pups. See figure S5e for changes to other proteins. Bottom: Bayesian network between measured variables, merged from networks inferred on DIV 6 and DIV 14 data. C) Cortical levels of excitatory synaptic proteins and their mRNA mouse pups at two ages, Log-FC in *Grin2a* +/-pups relative to litter-matched *Grin2a* +/+ pups. PF – prefrontal cortex, SS – somatosensory cortex. Error bars are mean ± s.e.m. across wells.

## Discussion

We present a multimodal procedure to acquire simultaneous information about live activity, biochemical processes and multiprotein compositions of neuronal synapses. Multimodal PRISM generates a fundamentally novel kind of data, namely large distributions of individual synapses in high-dimensional multimodal space. Analyzing these distributions by clustering, multivariate regressions, Bayesian network inference and conditional analyses enables structure-functional classification of synapses and reveals correspondence between classes, such as pre– and post-synaptically defined silent synapses. Network inference yielded predictive information about relationships between variables, whether they were driven by physical protein-protein or protein-mRNA interactions, reaction sequences or stoichiometric control.

### NMDAR antagonism plasticity and in vivo translatability

We applied this approach to study synaptic changes effected by ketamine and other NMDAR antagonists. Using combined multimodal imaging we identified, in a single experiment, excitatory synaptic potentiation in density, uneven protein composition, mRNA translation, and activity, providing molecular detail to the synaptogenesis that may generate ketamine’s antidepressant properties^18,65–68^. Bayesian network and conditional effect analyses not only retrodicted some of these phenotypes but also generated directly testable predictions about causal dependencies between these different synaptic changes. Based on similar observations in genetic Grin2a loss-of-function models and reduced Grin2a expression as seen with RIBOmap-PRISM, we hypothesize that the potentiating processes are a compensatory response to reduced presence of NMDARs in general or NR2A-containing NMDARs.

While many details are still unknown and rigorous orthogonal validations are still required, we identified the following processes:

1) Changes to synaptic activity including weaker glutamate release events and calcium levels immediately after drug removal combined with stronger potentiation over the following 1-2 hours. As these appear even in acute ketamine treatment and are necessary for APV-induced CaMKIIα potentiation, they likely precede the long-term molecular changes and may be mutually reinforcing with them.
2) Local synaptic increases in several proteins including AMPARs and NMDARs, scaffolding, cytoskeleton and active zone proteins. This is driven at least in part by TrkB activation and local protein synthesis at the synapse and strongly affects synaptic levels of NR2A.

The discrepancy between our observations and reports of TrkB pathway activation within much shorter time frames in vivo^68^ may stem either from differences between embryonic and postnatal neurons, possibly in NR2A/B ratios^75^, or differences in ketamine clearance. It may also be that the mutually reinforcing loop of increased synaptic activity to protein potentiation operates faster in a more densely interconnected 3D brain^76^. While in vitro cultures are considerably different from the specific neuronal wiring and interconnectedness resulting from brain development in vivo, we posit that they present a useful and faithful model of intrasynaptic relationships and drug-or gene-induced changes in synapse populations, when issues such as the above are accounted for to identify representative treatment regimes. Accordingly, we observed translation-based excitatory synaptic potentiation in vivo in Grin2a knockout mouse brains, as predicted by the in vitro measurements. The latter provide mechanistic details only accessible with Multimodal PRISM, demonstrating their value in supporting studies in animals.

### Caveats on interpretation

It is important to note certain limitations in interpreting data from different modalities. First, RIBOmap-PRISM reports on mRNA-protein associations, which can occur in the absence of translation, for example as static polysomes, including in synapses^40,77,78^. Second, Live-PRISM imaging occurs in tandem and not in parallel, so the resulting data combines protein and mRNA levels at the time of fixation with activity information from one hour leading up to fixation. Regression and conditional statistical analyses include confounds that must be accounted for. For example, the contribution of RIBOmap changes to the APV-induced effects on each protein in fig. 5c are likely exaggerated due to the small and non-representative fraction of synapses containing RIBOmap puncta. Finally, BN inference can be sensitive to data noise and thresholding of edge strengths, and hypotheses about causal connections between nodes generated by the model must be tested with direct perturbations as performed here and previously^49^.

### Generalizability and future outlook

Our modular approach, fundamentally requiring simple fluorescent optics, allows substitution of multiple forms of live imaging, including tracking movement/morphology, with activity integrators^79^ or reporters on membrane^80^ or other potentials, other neurotransmitters^81^, or psychedelic response^82^. It also allows integration of any biomolecular reporter that generates a DNA docking strand, including proximity ligation and ATAC-see^83^.

While focusing on synapses in this study, we expect our approach to be valuable for the study and phenotyping of populations of mitochondria, endosomes, phagosomes, and other organelles. The imaging method should also readily accommodate more biologically relevant samples such as tissue sections and iPSC-derived cultures. Finally, the high-dimensional multimodal synapse distribution that MM-PRISM generates provides access to rich phenotypes of drug treatments or genetic perturbations not accessible to other methods, such as marker correlations^49^, and that are potentially directly relevant to changes in neuronal circuit function. Thus, for example, application of this approach to human iPSC-derived neuronal cultures from patients living with schizophrenia or autism, for example, offers new hope for disease modeling and compound discovery in vitro.

## Supporting information

Supplemental Video 1 CAT-PRISM

Supplemental Video 2 GRL-PRISM

## Acknowledgements

This work was supported by NIMH R01-MH112694, NIMH R21-MH130624, and NSF-PoLS 1707999 awarded to M.B., as well as an Effective Ventures grant and the TVML foundation fellowship awarded to R.F. The work was also supported by the Center for Discovery of Therapeutics at the Broad Institute and use of the PerkinElmer Opera Phenix high-content/high-throughput imaging system funded by the S10 grant NIH OD-026839-01, as well as by MIT CEHS core center grant P30-ES002109 from the National Institute of Environmental Health Sciences. S.A. and M.S. were supported by the Stanley Center for Psychiaric Research. The authors would also like to thank Xiao Wang and Rena Ren for assistance with RIBOmap, Anna Lapteva for assistance with invader strand design, Opalina Vetrichelvan for Bayesian network-related calculations, Jasmine Wang for assistance with experiments, and Maria Alimova and Ranjith Muraleedharan for assistance with high-throughput imaging.

## Supplementary Information

**Supplementary Figure S1:**
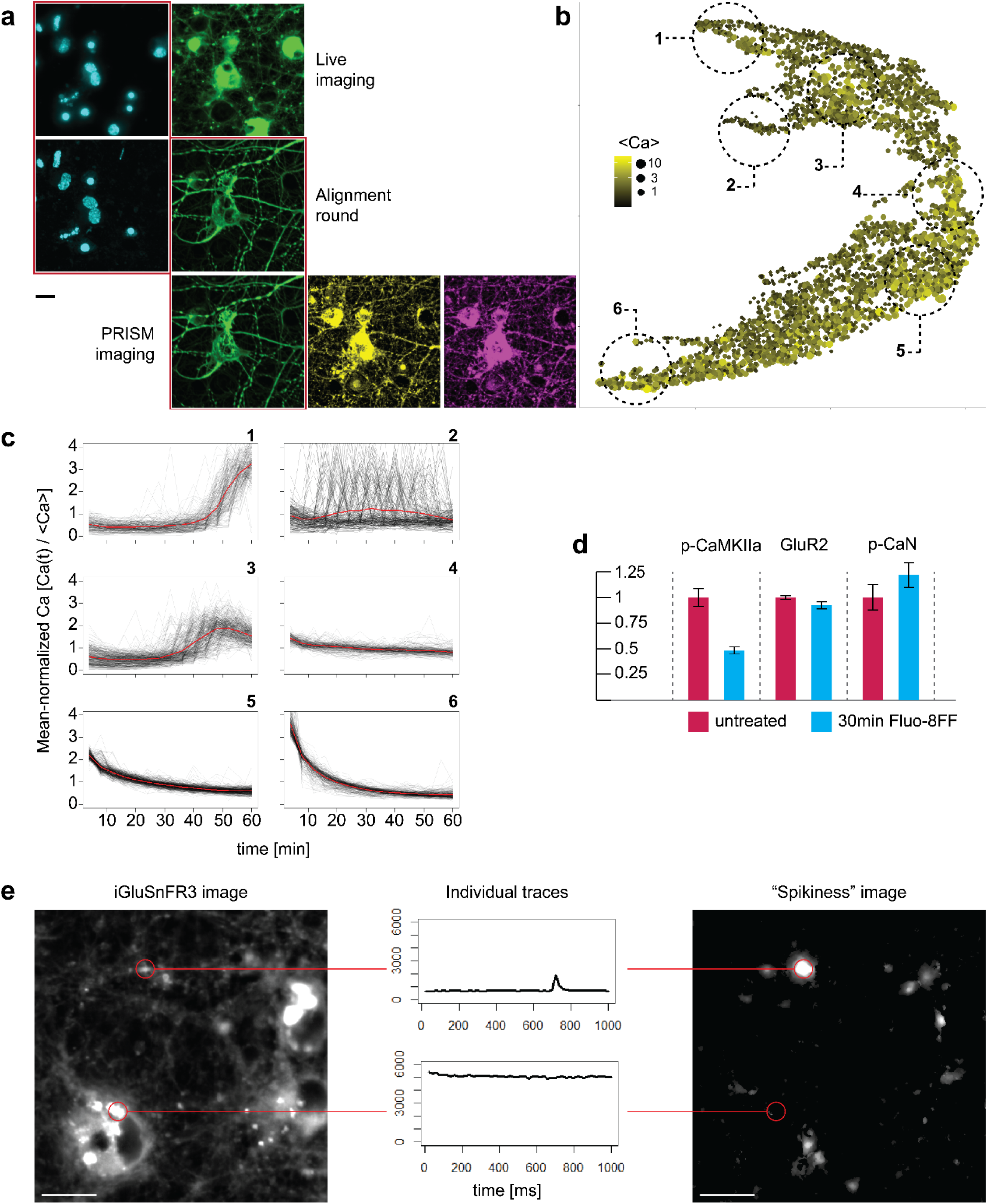
A) Live-PRISM alignment. Combined Hoechst and calcium imaging is aligned via the 405nm Hoechst channel to a post-fixation alignment round, which in turn is aligned to all PRISM imaging rounds via the 488nm MAP2 channel. B) UMAP of synaptic calcium trace shapes (from distances assigned by DTW), colored and sized by average calcium signal. C) Individual (gray) and average (red) calcium traces in regions 1-6 circled in (B). D) Average synaptic levels of phospho-CaMKIIα, GluR2, and phospho-Calcineurin, in neurons exposed to 10uM Fluo-8FF. E) Maximum projection across a representative exposure window in iGluSnFR3 imaging and its corresponding spike analysis image, with traces from a weak but spiking and a strong but nonspiking region highlighted. A,F scale bars are 10um.

**Supplementary Figure S2:**
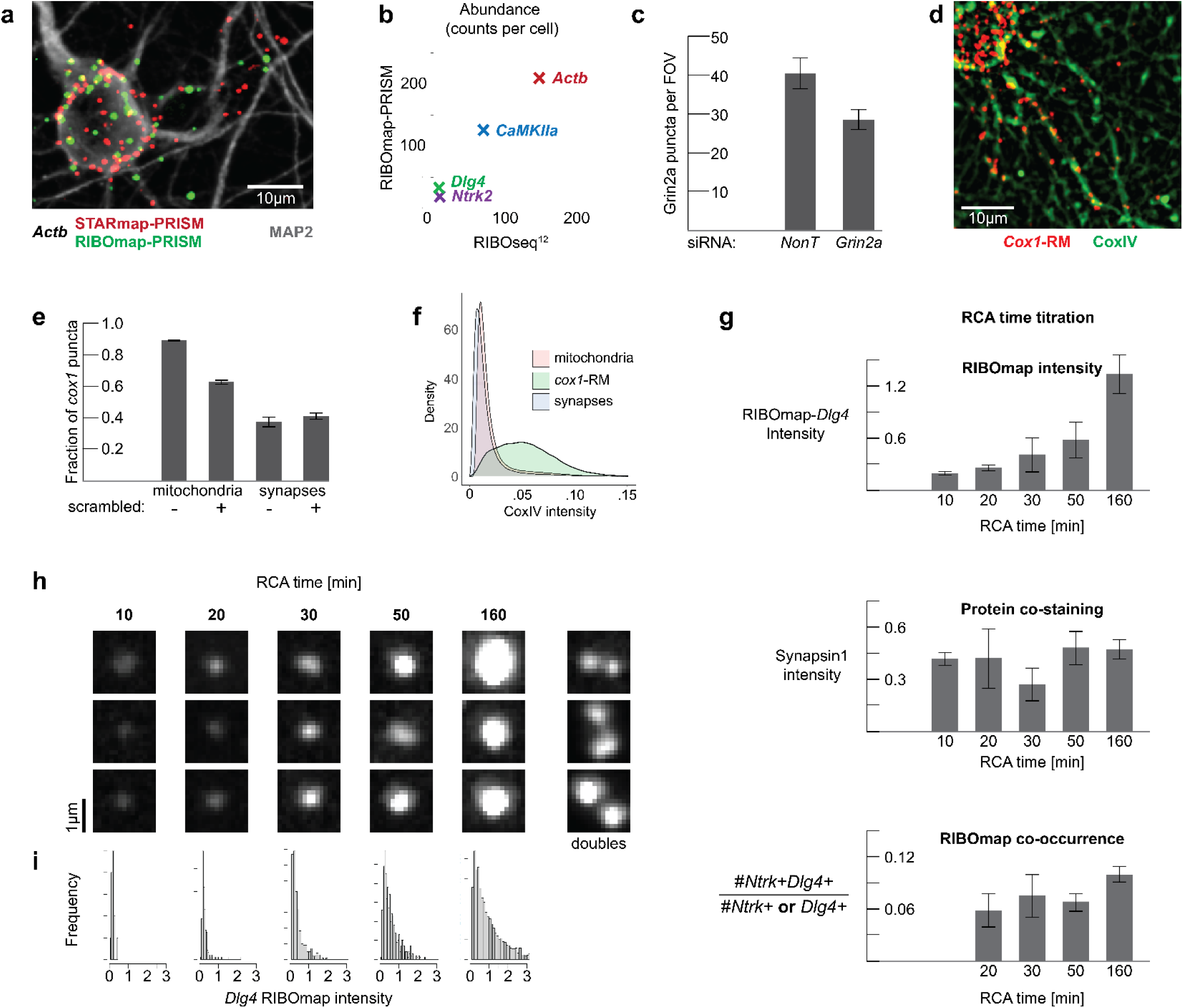
RIBOmap-PRISM and STARmap-PRISM. A) STARmap-PRISM puncta (red) and RIBOmap-PRISM puncta (green) of *Actb* costained with MAP2 (white). Scale bar is 10um. B) Estimates of puncta count per cell count (Methods). Y-axis; this study. X-axis; Glock et al. C) Total RIBOmap-PRISM puncta counts per field of view of *Grin2a* after treatment with siRNA mix or nontargeting siRNA (NonT). D) RIBOmap-PRISM puncta of cox1 overlaid with staining for CoxIV used to identify mitochondria. E) Fraction of cox1 puncta that overlapped with CoxIV-identified mitochondria or Bassoon-identified synapses, in aligned (-) and rotated (+) images. F) Histograms of average CoxIV intensity in three sets of objections – mitochondria (red), synapses (blue), and cox1 RIBOmap puncta (green). G) Top: Average *Dlg4* RIBOmap-PRISM intensity integral in synapses positive for *Dlg4*, in cultures that underwent different amplification times in the RCA step. Middle: Average Synapsin1 signal in *Dlg4*-positive synapses. Bottom: Fraction of puncta which are positive for both *Ntrk2* and *Dlg4*, out puncta positive for either. The 10-minute amplification time showed too few *Ntrk2*-positive puncta for a reliable estimate. H) Representative RIBOmap-PRISM puncta from different amplification times. Right column shows pairs of adjacent puncta assigned to the same synapses which were segmented properly. I) RIBOmap-PRISM intensity histograms of *Dlg4* puncta in different amplification times.

**Supplementary figure S3:**
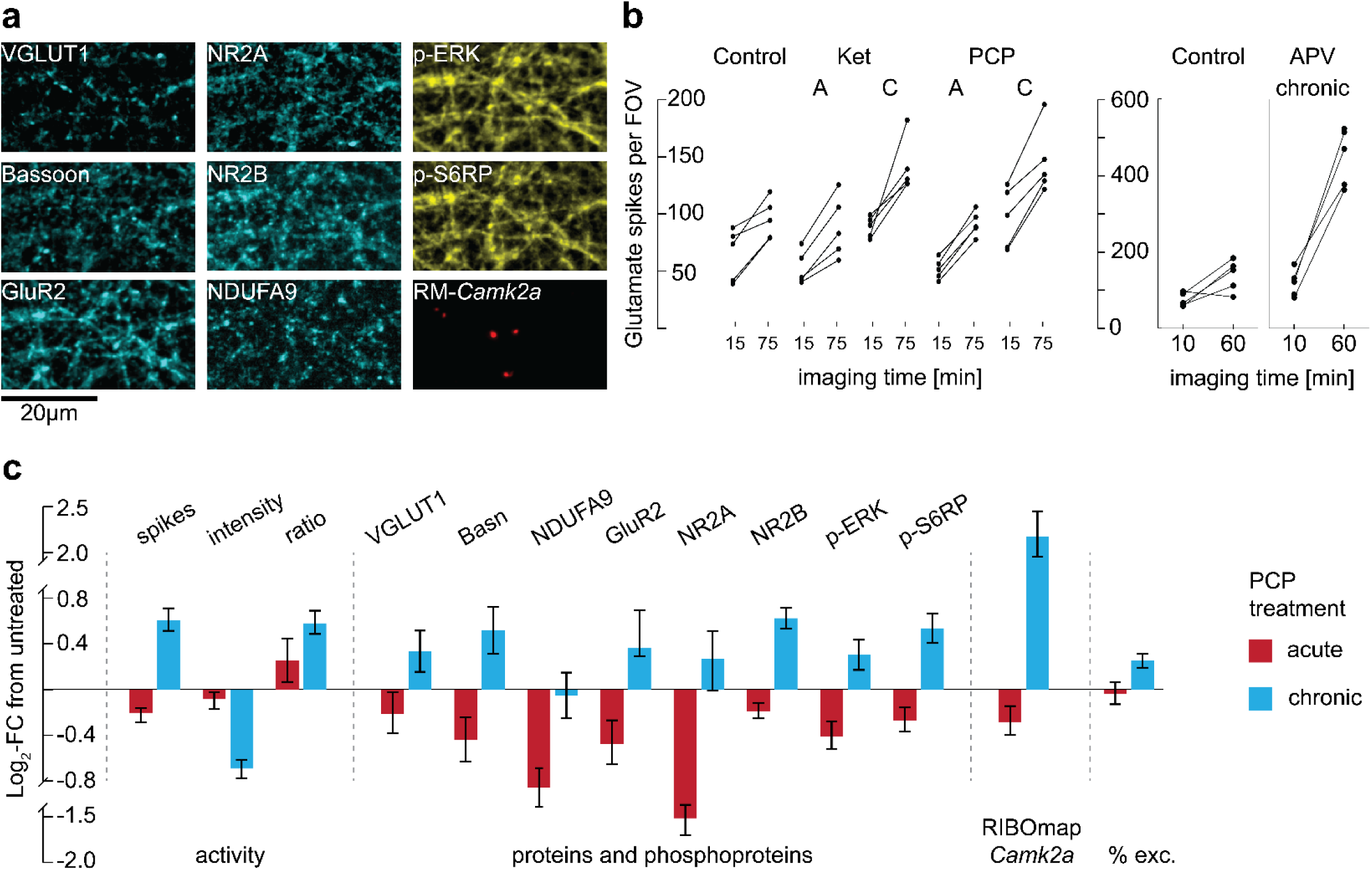
A) Aligned images of different proteins, phosphoproteins and RIBOmap-PRISM puncta in the same field of view as Fig. 3b. B) Total glutamate spike count per FOV in each culture at the start and end of live imaging, across treatment groups (Ketamine, PCP and APV) and regimes (acute, “A” or chronic, “C”). C) As figure 3c-e but under PCP treatment.

**Supplementary figure S4:**
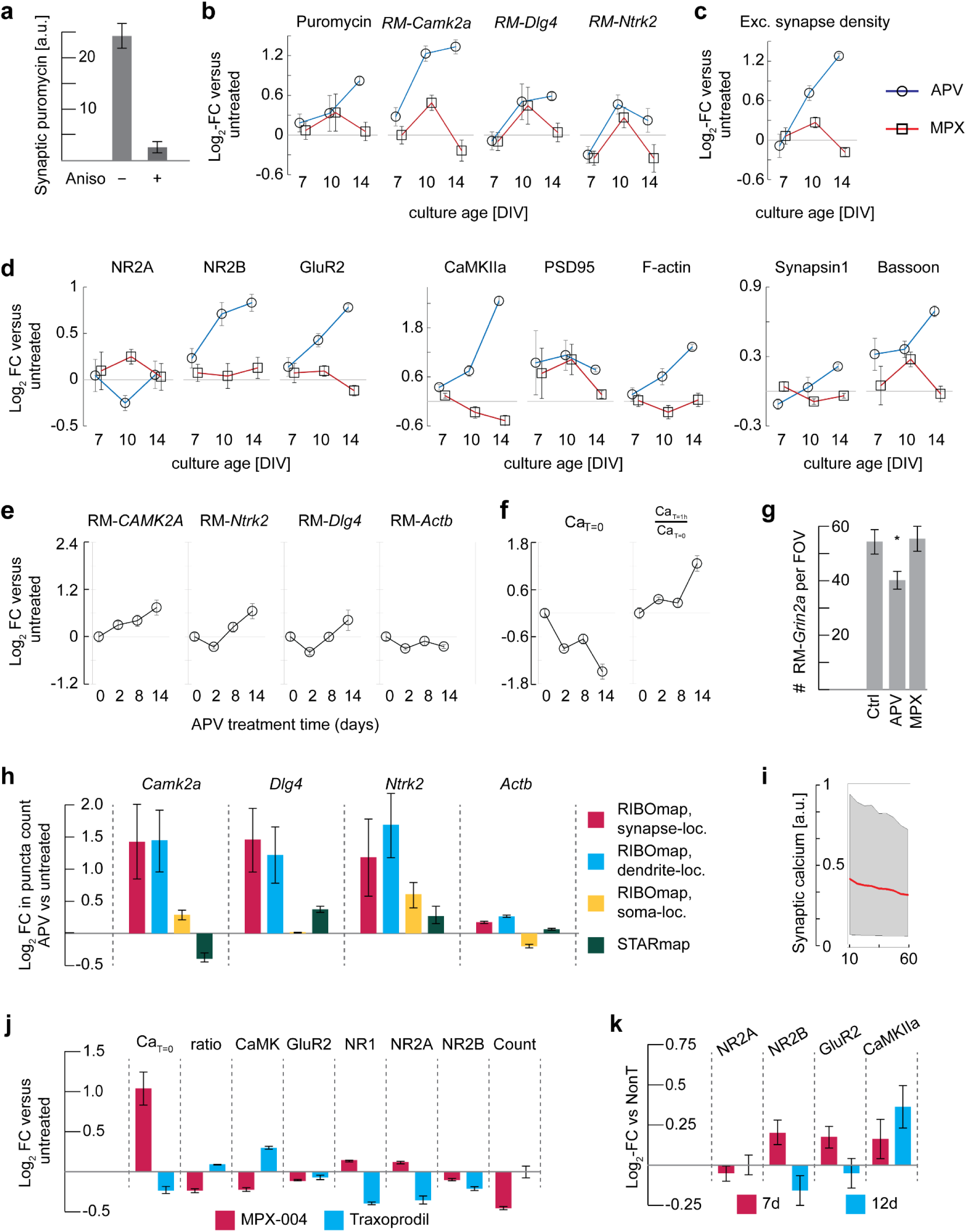
A) Average synaptic puromycin levels with and without concomitant addition of anisomycin. B-D) Log_2_-fold changes to synapses in cultures treated at DIV 5 with APV (circles, blue line) or MPX (squares, red line) and measured at DIV 7, 10 or 14, relative to untreated cultures of the same age. B) Average synaptic levels of puromycin and presence of RIBOmap-PRISM puncta presence of *Camk2a*, *Dlg4* and *Ntrk2*. C) Density of excitatory synapses. D) Average synaptic levels of eight proteins, grouped by type. E-F) Log_2_-fold changes to synapses in DIV 19 cultures treated with APV for 2, 8 and 14 days before measurement, relative to untreated cultures (“0”). E) RIBOmap-PRISM puncta presence of four genes, F) Starting calcium and calcium ratio over an hour of measurement after replacement with APV-free media. G) Total number of RIBOmap-PRISM puncta of *Grin2a* per field-of-view in untreated and APV and MPX-treated cultures. H) Log_2_-fold changes in total RIBOmap– and STARmap-PRISM (green) puncta counts of four genes in APV-treated vs untreated cultures, grouped by colocalization with Synapsin1/MAP2-identified neuronal structures – synapses (red), dendrites (blue), soma (yellow). I) Like fig 3b, average (red) and 5-95^th^ percentile calcium traces (gray) in MPX-004-treated cultures. J) Head-to-head comparison of chronic treatment with MPX vs Traxoprodil, Log_2_-fold changes to synaptic calcium starting levels, ratios over one hour, five proteins, and excitatory synapse count, versus untreated cultures. K) Log_2_-fold changes in average synaptic protein levels in cultures treated with siRNA against *Grin2a* vs Nontargeting, at DIV 6 (blue) and 11 (red) before fixing on DIV 18.

**Supplementary Figure S5:**
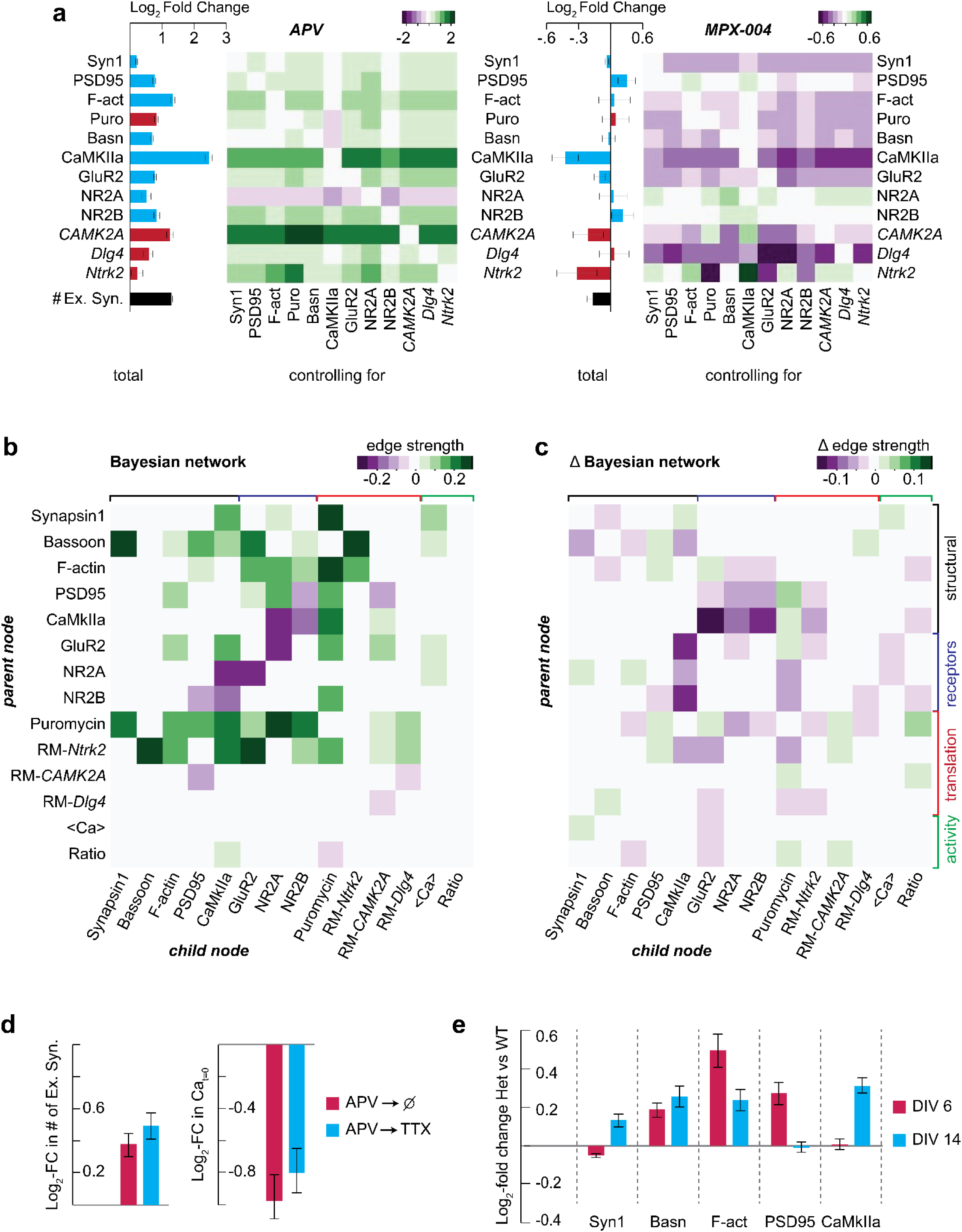
A) Log_2_-fold changes in APV-(left) and MPX-(right) treated cultures, in proteins (blue), translation indicators (red) and overall synapse density (black), relative to untreated cultures. Bars: Overall changes. Heatmaps: Changes in each variable when controlling for each of the other variables by stratification (see methods). B) Full Bayesian network of eight proteins, four translation indicators, and two activity indicators, in untreated cultures, presented as a heatmap of edges from parent node (rows) to child node (columns). Green denotes positive edges, purple denotes negative edges. C) Changes to the strengths of edges in B in APV-treated cultures. D) Log_2_-fold changes, relative to untreated cultures, in cultures chronically treated with APV then replaced with either imaging media alone (red) or imaging media + 2uM TTX (red). Left: Excitatory synapse density, Right: Starting calcium levels. E) Log-FC to average levels of other synaptic proteins apart from those in fig. 5b in *Grin2a*+/-mouse neurons relative to *Grin2a*+/+. Error bars are mean +/-s.e.m. across wells.

## Methods

### Orthogonal imaging-docking sequence pair generation

Sequences 1-12 were identical to those previously published^42^ except LNA imaging strands were extended with a 5’ four-nucleotide toehold (ACTC) for toehold-mediated strand displacement by invader strands (see below). Additional sequences (13-29) were generated by screening a randomly generated library of 11-nt docking strands with GC content between 10 and 30% against existing LNA imaging strands using our previously published algorithm for crosstalk prediction^42^. Lowest scoring sequences were converted to 15-nt LNA imaging strands with all possible variations on locked base positions and screened against all existing and new DNA docking strands. This generated 17 docking-imaging pairs with low predicted crosstalk scores against each other and published sequences. Crosstalk was tested experimentally by binding biotinylated docking strands to a streptavidin-coated plate (ThermoFisher) and measuring fluorescent LNA signal after probe binding and washing as described below. Two sequences were removed after exhibiting some crosstalk with existing sequences. Another sequence was removed after testing for nonspecific binding to unstained, RNase-treated neurons. This resulted in 14 new sequences of which seven were used alongside previously published ones.

### Imaging reagent preparation

Antibody-DNA conjugates were prepared with either chemical (SMCC) or enzymatic (SiteClick) conjugation as described previously. Table S1 details the antibodies used for multiplexed imaging, including the conjugation strategy used for each. Table S2 details sequences of docking strands, imaging strands, invader strands, and RIBOmap probes.

Chemical conjugation was done using 4-(N-Maleimidomethyl)cyclohexane-1-carboxylic acid N-hydroxysuccinimide (SMCC), a heterobifunctional molecule which binds exposed amines on the antibody via an N-hydroxysuccinimide (NHS) moiety on one end and binds thiol-oligonucleotides via a maleimide on the other end. Antibodies were purchased as formulations without serum proteins as listed below (Table S1), 0.1-1 mg of antibody were purified into PBS and concentrated to 1 mg/ml using Amicon Ultra centrifugal filters (10 kDa, EMD Millipore). From a freshly prepared stock at 2 mM in DMF, SMCC (Sigma Aldrich) was added to the antibody at 7.5x molar excess. The reaction mixture was protected from light and incubated for 3 h at 4°C on a shaker. Excess SMCC was removed by purification into PBS using the above Amicon centrifugal filters.

In parallel, 25 nmol 5’ thiol-modified ssDNA (Integrated DNA Technologies, modification catalog no. /5-ThioMC6-D/) were deprotected by mixing with 150mM DTT in 0.5x PBS, protected from light and incubated for 2 h at 25°C on shaker. The reduced 5′ thiol-modified ssDNA was purified into water using NAP-5 columns (GE Life Sciences). The reduced 5’ thiol-modified ssDNA was spin vacuum dried, resuspended in 50ul water and added to the antibody-SMCC conjugate at 15x molar excess, the reaction mixture was protected from light and incubated overnight at 4°C on a shaker. Antibody-ssDNA conjugates were purified into PBS using Amicon Ultra centrifugal filters (10 kDa, EMD Millipore). Amino-modified phalloidin (Bachem) was conjugated using the procedure described above, but with the following changes: the molar excess of SMCC was 10x and the molar excess of reduced 5’ thiol-modified ssDNA was 1x. HPLC purification was employed to remove unreacted SMCC and 5’ thiol-modified ssDNA, respectively (Waters, BEH C18 column, gradient for phalloidin-SMCC: from 80% TFA in water and 20% acetonitrile to 20% TFA in water and 80% acetonitrile over 10 min, gradient for phalloidin-ssDNA: from 90% 0.1 M TEAA in water and 10% acetonitrile to 60% 0.1 M TEAA in water and 40% acetonitrile over 10 min). Antibody concentrations were determined by absorbance measurements at 280 nm.

Enzymatic conjugation was done using the SiteClick™ (Invitrogen) conjugation technique following the manufacturer’s protocol. This technique replaces the Fc galactoses on the antibody with azide-modified sugars, which then react with a DBCO-modified oligonucleotide. 0.1-0.2mg of antibody are concentrated to 1-2 mg/ml in 1x Tris buffer and incubated with β-galactosidase. Azide-modified, terminal galactosides were attached using β-galactosyltransferase. Azide-modified antibody was purified into 1x Tris buffer using Amicon Ultra centrifugal filters (50 kDa, 4000 g, EMD Millipore).

In parallel, DBCO-modified docking strands were prepared by mixing 25nmol 5’ amine-modified ssDNA (IDT) with a 20-fold excess of DBCO-NHS (Sigma-Aldrich) from a DMF solution, at 1:1 mix of DMSO and and PBS, for 3 hours at 25C. Excess DBCO was removed by 2x purification on NAP-5 columns and the conjugate was spin vacuum dried, resuspended in 50ul water and added to the azido-modified antibody at a 20-fold excess, then incubated for 48 hours at 4C. The conjugates were then purified from excess docking strand by repeated concentrations on 30k MWCO Amicon centrifugal filter until the 260/280 ratio was stable. All antibody-ssDNA conjugates were stored at –20°C in PBS with 50% glycerol.

Imager strands: 5 nmol of 5’/3’ diamino-modified ssLNA (IDT) was mixed 100 nmol of NHS-Alexa Fluor 568 or NHS-AlexaFluor 647 (Sigma Aldrich) in 1:1 PBS:DMSO. Following immediate vortexing, the reaction mixture was protected from light and incubated overnight at 25°C on a shaker. Excess dye was removed using NAP-5 columns (GE Life Sciences). Fractions containing ssDNA were identified using absorbance measurements at 260 nm. Subsequently, 0.1 M TEAA was added ssLNA-dye conjugates and conjugates bearing two dyes were purified by HPLC (Waters, BEH C18 column, gradient for Atto 565: from 80% 0.1 M TEAA in water and 20% acetonitrile to 70% 0.1 M TEAA and 30% acetonitrile over 10 min, gradient for Atto 655: from 90% 0.1 M TEAA in water and 10% acetonitrile to 75% 0.1 M TEAA in water and 25% acetonitrile over 10 min). Peaks corresponding to ssLNA conjugates bearing two, one or no dye were assigned based on absorbance spectra. Solvents were removed in vacuo and ssLNA-dye conjugates were dissolved in water to 5 to 20 μM, depending on the yield.

### Rat embryonic neuron culturing and treatment

For all experiments except the Grin2a KO experiment, we used neurons from E18 embryonic rat hippocampus dissociated and plates as described previously. Hippocampi from embryos in separate pregnant Sprague Dawley rats were supplied by BrainBits LLC in Hibernate E, digested in Hibernate E containing 20 U/ml Papain (Worthington Biochem), and 0.01% DNase (Sigma-Aldrich) for 10 min at 37C and triturated. Neurons were spun down at 210g for 2×5min, cell pellets were then resuspended into NbActiv1+25uM glutamate (BrainBits LLC, now TransnetYX), and plated at a density of 22,000 cells/well onto poly-d-lysine-coated, black-walled, thin-bottomed 96-well plates (Corning BioCoat). After 24 hours, AraC was added to each culture at a concentration of 1uM, to suppress glia proliferation and minimize well-to-well variability resulting from it. At DIV 5 (or DIV 4 when drug treatment was started on DIV 5), the media was entirely replaced with preequlibrated NbActiv4 (BrainBits LLC) and subsequently maintained in this media. In experiments that involved puromycin labeling, 250nM puromycin (Sigma Aldrich) were added 15min before fixation. For anisomycin controls, puromycin labeling was performed after 30 minute pre-treatment of neurons with anisomycin (final concentration of 10 µg/ml).

All drug reagents were purchased from Sigma Aldrich. For the treatments in figure 2, neurons were treated with either 1µM TTX, 50µM APV, 20µM bicuculline or 20µM bicuculline + 200µM glycine in BrainPhys media immediately before live imaging. Accell self-transfecting siRNA mixes against Dlg4 and Grin2a were purchased from Horizon Discovery, and added at 1µM to cultures at DIV 6. Ketamine and PCP were purchased from Sigma Aldrich and added at DIV 6 or 14 at concentrations of 10µM and 5µM, respectively.

### Grin2a knockout neuron culturing

Female mice heterozygous for Grin2a (B6;129S-Grin2a<tm1nak>, RIKEN BioResource Research Center) were paired with male wild-type (C57BL/6J, Jackson Labs 000664) mice. On embryonic day 16.5, a pregnant dam was euthanized by CO2 inhalation in accordance with the Institutional Animal Care and Use Committee guidelines. Pups (n=8) were surgically retrieved and the cortex of each pup was dissected out into ice-cold HBSS supplemented with glucose. The embryonic tissue was digested in 0.125% trypsin for 15 minutes (37 °C), triturated in ice-cold HBSS with a fire-polished glass pipette, and the resulting cell suspension passed through a 40 um filter. Cells from each pup were counted and plated on a black-walled clear-bottom poly-D-lysine coated 96-well plate (Corning) at a density of 20,000 cells/well (6 replicates per pup). Neurons were cultured in NbActiv4 medium (TransNetyx LLC) supplemented with penicillin/streptomycin in a 5% CO2 incubator at 37 °C. Cytosine arabinoside (Ara-C; 1 µM final concentration) was added at DIV1; media with Ara-C was discarded on DIV2 and fully replaced with fresh media. Neurons were supplemented with additional fresh media every 3-4 days.

### CAT-PRISM

Neurons were treated at T-45min before imaging with 5uM Fluo-8FF (AAT Bioquest) and at T-10min with 10uM Hoechst 33342. At T=0, the media was replaced with pre-equilibrated BrainPhys Imaging Medium^84^ (StemCell Technologies) containing any indicated chemical treatment and imaging started at approximately T+10min. Calcium imaging was performed as repeated minimal exposure confocal imaging in the 488 channel over the entire plate, so that each field of view was imaged once every 4-6 minutes over an hour. At the end, a single round of confocal imaging in the 405 and 488 channels was performed, followed by immediate fixation with PFA.

### GRL-PRISM

pAAV.hSyn.iGluSnFR3.v857.GPI was a gift from Kaspar Podgorski (Addgene viral prep # 178331-AAV1). At DIV 9 and until imaging, neurons were transduced with AAV1 vectors carrying iGluSnFR3.v857.GPI^46^ at 10^5^ MOI. Prior to imaging, neurons were treated with 10uM Hoechst for 10min then replaced with pre-equilibrated BrainPhys Imaging Medium^84^ containing any indicated chemical treatment. Glutamate imaging was done in widefield (epifluorescence) format in bursts of 50-100 20ms exposure frames in the 488 channel and minimal power, leading to 1-2s recording windows at 50Hz. To measure spiking potentiation, the same fields of view were imaged twice, 1hr apart. After glutamate imaging, a single round of confocal imaging in the 405 and 488 channels was performed, followed by immediate fixation with PFA.

### RIBOmap-PRISM and STARmap-PRISM

RIBOmap-PRISM and STARmap-PRISM primer and padlock probe sequences were generated the following way: First, 50-60nt unique hybridization regions against the coding region of the mRNA transcript of interest were identified that have minimal sequence similarity to other transcripts expressed in neurons. These regions – approximately 10-25 per target gene – were split into primer and padlock hybridization regions of 19-26nt, Tm∼60C, with one nucleotide separation. Primer and padlock hybridization sequences were inserted into RIBOmap probe sequences from Zeng et al^47^. In addition, the 6-nt barcode sequences in the padlock probes were replaced with 11-nt PRISM docking sequences. The resulting full primer and padlock sequences were screened against both the rat transcriptome and against the full set of DNA-LNA probe pairs (as described above) to pick 3-5 probe pairs for each transcript of interest that have minimal crosstalk.

RIBOmap staining was performed similar to the protocol by Zeng et al. ^47^, with some modifications. Cultures were fixed with PFA+sucrose then permeabilized with PBS containing 0.25% Triton X-100 and 2mM Vanadyl Ribonucleoside Complex (VRC, Sigma Aldrich) for 10min at room temperature. After washing with PBS+0.1% Tween-20 (PBST), cultures were incubated with the hybridization solution overnight at 40C with shaking. Washing, ligation and amplification were performed as described by Zeng et al., except that amplification was performed for 30 minutes in most experiments and 10-160 minutes in the experiment in figure S2. Following amplification, the cultures were fixed again with 2% PFA for 10min, after which they underwent the standard PRISM protocols. STARmap staining was performed identically but ribosome splint probes were excluded during the hybridization step.

### Protein-PRISM staining

Staining was performed as described previously^42,44,49,85^. Fixed cultures were permeabilized with 0.25% Triton X-100 (Sigma Aldrich) for 10min, underwent RNase treatment at 37C for 1hr, then blocked with 5%BSA (Sigma Aldrich) + 1mg/mL ssDNA (ThermoFisher) in PBS, which was also the composition for all subsequent antibody solutions. Antibody staining was performed in three steps – overnight staining with primary antibodies against targets that were imaged with secondary PRISM, then secondary staining with conjugated secondary antibodies (as well as a 488-labeled secondary antibody for the MAP2 staining), then another overnight staining round with conjugated primary antibodies. After rounds of antibody staining neurons were post-fixed with 2% PFA for 10min, then stained with 10uM Hoechst and washed with imaging buffer.

### Protein– and RIBOmap-PRISM imaging

With Live-PRISM imaging, an alignment round was done imaging Hoechst and MAP2 staining simultaneously in all fields previously live imaged. Then, in every PRISM imaging round, the desired AF568– and AF647-labeled LNA imaging probes were introduced at 20nM for 10min in imaging buffer (PBS+0.5M NaCl), washed with imaging buffer and the neurons imaged in the 488, 568 and 647 channels. After each round of imaging, probes were removed by 10min incubation with 4uM invader strands in imaging buffer, followed by washes in 37C wash buffer (0.01x PBS). Before introducing the next LNA imaging probe pair, the plate was imaged again to ensure removal of the 568 and 647 signals. In experiments that imaged NR2B or Tim23 these targets were imaged in the last round with an AF568-labeled secondary antibody against Mouse IgG2b, alongside nuclear imaging with DAPI or imaging of 405-labeled probes.

### Imaging setup

All imaging was done using an Opera Phenix high content spinning disk confocal microscope (PerkinElmer) at the Broad Institute CDoT. Imaging was done with a 63x water immersion objective, collecting on a 2160×2160 detector (6.5um pixel size) with 2×2 binning for 1080×1080 raw images at 0.18um/px resolution. Z-stacks of 4-7 planes (steps of 1-2um) were collected in confocal imaging. Live imaging was performed at 37C with 5% CO2.

### Image processing – CAT– and Protein-PRISM and IF images

All images from calcium live, PRISM and regular immunofluorescence imaging underwent the following processing steps before alignment and analysis: 1) Calculate of an illumination profile for each channel by calculating by-pixel medians across all images of that channel, and divide the images by the illumination profile. 2) Calculate background intensities for each image consisting of the average of the 5% lowest-intensity pixels in 30×30px (for images of synapses), 60×60px (for MAP2) or 120×120px (for nuclear staining) windows and subtracting from each illumination corrected image its background image. 3) Maximum projection of images across Z-stacks.

### Image processing – GRL-PRISM

Image sets from this modality underwent illumination correction and background subtraction as described above, followed by calculation of an average bleaching curve used to correct videos for photobleaching. The average of each image set was subtracted from all images of that set, and the resulting set underwent median filtering with a 5px window. The resulting filtered set was then fed into a by-pixel spike detection algorithm that labels individual pixels by the quantity and total intensity of spikes (values sufficiently higher than surrounding values) in the time plot for that pixel. All image processing was done in MATLAB with bespoke scripts (supplementary code).

### Organelle segmentation and quantification

Images processed as described above were aligned across live imaging rounds, PRISM imaging rounds and between modalities, cropped, and analyzed using a bespoke CellProfiler pipeline (supplementary code) with the following core steps:

1) The DAPI image is used to identify nuclei objects. All other images of the same field are then masked by the nuclei to prevent artifacts from non-specific nuclear localization of the antibodies.
2) The MAP2 image is used to identify dendrite objects.
3) When calcium imaging was done, a maximum projection of images across the timelapse was used to extract calcium puncta. When glutamate imaging was done, the spike detection algorithm output image was used to extract glutamate spiking puncta. For all antibody-PRISM and RIBOmap-PRISM images the maximum projection image for that channel was used to extract puncta of that protein or transcript.
4) A white top hat filter with a radius of 6px is applied to all synaptic protein images across all rounds to enhance puncta.
5) For initial synapse counting analysis synaptic objects were segmented and identified in images of each channel by applying the RobustBackground tool, which calculates an optimal threshold value for each 20px window individually based on the intensity histogram. When comparing treatment groups, we used a global threshold derived from the average threshold calculated by RobustBackground across all imaged fields. We then applied this value as a uniform threshold to all images of that channel to ensure that all images are segmented identically.
6) Synapsin1 puncta are then masked using the dendrites previously identified, to retain only puncta which are within 12px of a dendrite. These are then defined as synapses.
7) Puncta in all other channels are assigned to synapses if they overlapped with Synapsin1 puncta more than 6.25% (for postsynaptic proteins) or more than 50% (for presynaptic proteins).
8) Finally, levels of each marker – protein, phosphoprotein, ribosome-bound transcript, calcium signal at each individual time point, and glutamate spike count and intensity – per synapse are calculated as the intensity integral of that protein’s image across its punctum. If a certain marker did not have an identified puncta associated with a synapse, its level was marked as 0.

### Soma ID and cell body counting

Extrasynaptic neuronal soma regions were identified from MAP2 images as described before ^49^. Images underwent a tophat filter with a 60px kernel size which mostly removed dendrite regions and retained only non-dendrite MAP2 staining. The resulting image was then masked for nuclei and synapses from DAPI and Synapsin1 staining, respectively.

Estimate of ribosome-associated transcripts per cell for RIBOmap-PRISM was done by dividing total number of counts to total number of ID’d soma. Estimate of ribosome-associated transcripts per cell for Ribo-Seq was done using the raw counts reported by Glock et al.^1^ from 200ng mRNA and estimating 30pg mRNA per neuron.

### CAT-PRISM analysis

DTW was performed on mean-normalized calcium traces associated with Synapsin1 puncta using the dtw package in R^57^. This generated a distance matrix of calcium trace similarity. For each pair of calcium-positive synapses, the trace similarity distance was combined with the Euclidean distance between protein and mean calcium measurements to generate a distance matrix in combined activity-multiprotein space, which was then visualized using a UMAP projection with the umap package in R.

### Bayesian network analysis and controlled edge and treatment effect calculation

Prior to Bayesian network inference on synapses, data was filtered to include only excitatory synapses (negative for Gephyrin and/or vGAT, when measured, and positive for at least one excitatory synaptic protein out of PSD95, CaMKIIa, phospho-CaMKIIa, GluR2, NR2A, NR2B, VGLUT1).

Data was discretized into 20 bins (with 0 values in a separate bin) and network derivation was done using the likelihood-score-maximizing ‘tabu’ algorithm6 and boot.strength functions in the R package bnlearn^86^, which performed 50 repeats of network inferences from resamplings of 10% of the data. This yields both the fraction of repeats in which a certain network edge appeared and the fraction of appearances in which the edge was in a specific direction.

Given a network, we define the strength of an edge between two nodes as the average correlation of the two variables across strata where the other parents of the daughter node are held constant^87,88^. That is, the strength of an edge from A to B, where B also has edges leading to it from n other variables, for example C and D with n=2, as the correlation between A and B when controlling for C and D. To estimate that, we repeated the following algorithm to calculate average correlations between A and B across strata of equal C and D:

– Sample a point (A_0_, B_0_, C_0_, D_0_)
– Find set of all points (A, B, C, D) such that 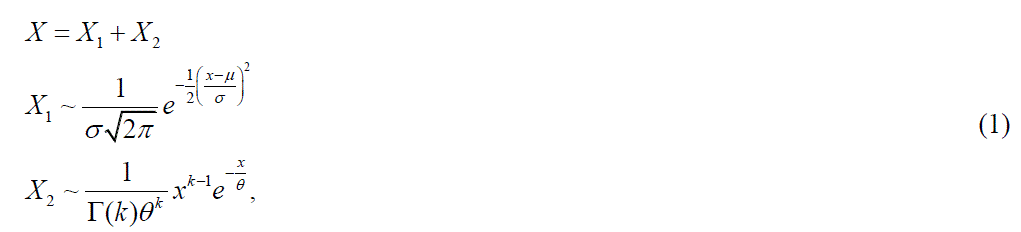 where n is the number of variables to control for (2 in this example) and ε is a predetermined tolerance level set at 0.5 (smaller tolerances did not yield significantly different measures)
– If the set contains more than 5 points, calculate Pearson’s correlation coefficient *cor* (*A, B*) across that set.
– Average the resulting correlation measure across 20 · 2^*n*^ such samplings.

A similar stratification procedure was done to assess the conditional effect of a certain treatment on protein A when controlling for proteins B and C. Treated and controls groups were pooled together, a point was sampled at random and a set of all points with similar B and C was found, and the log_2_-fold difference between the mean levels of A in treated vs control synapses was calculated and averaged across many samplings.

Unless indicated otherwise, the Bayesian networks presented include only those edges which appeared in >80% of bootstrapped repeats, edge directionality was given when this directionality appeared in >60% of bootstrapped repeats. For Bayesian network inference in Grin2a knockout mouse neurons, inference was performed separately on DIV 6 and DIV 14 data. The edges presented are only those which appeared in the networks from both datasets with edge strengths (controlled correlations) above 0.1 in both.

### RNA-seq in Grin2a knockout mice

RNA-seq was performed as described previously^89^. RNA was prepared from micro-dissected tissues using RNeasy Mini Kit (Qiagen) following the manufacturer’s instructions. Briefly, tissue samples were lysed and homogenized, placed into columns, and bound to the RNeasy silica membrane. Next, contaminants were washed away, columns were treated with DNase (Qiagen) to digest residual DNA, and concentrated RNA was eluted in water. RNA concentration was measured using a NanoDrop Spectrophotometer and RNA integrity (RIN) was measured with RNA pico chips (Agilent) using a 2100 Bioanalyzer Instrument (Agilent). Purified RNA was then stored at –80°C until library preparation for bulk RNA-seq analysis. Bulk sequencing libraries were prepared using a TruSeq Stranded mRNA Kit (Illumina) following the manufacturer’s instructions. 200 ng of isolated total RNA from each sample was used and the concentration of resulting cDNA library was measured with High Sensitivity DNA chips (Agilent) using a 2100 Bioanalyzer Instrument (Agilent). A 10 nM normalized library was pooled, and sequencing was performed on a NovaSeq S2 (Illumina).

### Postsynaptic density proteomics in Grin2a knockout mice

Protein extraction and quantification was performed as described previously^90^. Flash-frozen DLPFC gray matter (human) or whole cortex (mice) was dounce-homogenized in ice-cold homogenization buffer (5 mM HEPES pH 7.4, 1 mM MgCl2, 0.5 mM CaCl2, supplemented with phosphatase and protease inhibitors). The homogenate was centrifuged for 10 minutes at 1,400 g (4°C) and the supernatant was re-centrifuged at 13,800 g for 10 minutes (4°C). The resulting pellet was resuspended in 0.32 M Sucrose, 6mM Tris-HCl (pH 7.5) and layered gently on a 0.85 M, 1 M, 1.2 M discontinuous sucrose gradient (in 6mM Tris-HCl pH 7.5) and ultracentrifuged at 82,500 g for 2 hours (4°C). The synaptosome fraction—which sediments at the 1 M and 1.2 M sucrose interface—was collected, an equal volume of ice-cold 1% Triton X-100 (in 6mM Tris-HCl pH 7.5) was added, mixed thoroughly and incubated on ice for 15 minutes. The mixture was ultracentrifuged at 32,800 g for 20 minutes (4°C), and the final synapse protein pellet was collected by resuspension in 1% SDS. A small aliquot was taken to measure the protein concentration using the BCA assay (ThermoFisher Scientific), and the remaining protein was stored at –80°C until being processed. Protein from purified synaptic fractions was digested as described above. 100 μg of each sample was labeled with a TMT16 reagent following the protocol described above. The combined sample was fractionated on a 4.6 mm x 250 mm Zorbax 300 extend-c18 column (Agilent) and concatenated into 18 fractions for LC-MS/MS analysis.

